# Decoding WW Domain Tandem-mediated Target Recognitions in Tissue Growth and Cell Polarity

**DOI:** 10.1101/694976

**Authors:** Zhijie Lin, Zhou Yang, Ruiling Xie, Zeyang Ji, Kunliang Guan, Mingjie Zhang

## Abstract

WW domain tandem-containing proteins such as KIBRA, YAP, and MAGI play critical roles in cell growth and polarity via binding to and positioning target proteins in specific subcellular regions. An immense disparity exists between promiscuity of WW domain-mediated target bindings and specific roles of WW domain scaffold proteins in cell growth regulation. Here, we discovered that WW domain tandems of KIBRA and MAGI, but not YAP, bind to specific target proteins with extremely high affinity and exquisite specificity. Via systematic structural biology and biochemistry approaches, we decoded the target binding rules of WW domain tandems from cell growth regulatory proteins and uncovered a list of previously unknown WW tandem binding proteins such as β-Dystroglycan, JCAD, and PTPN21. The WW tandem-mediated target recognition mechanisms elucidated here can guide functional studies of WW domain proteins in cell growth and polarity as well as in other cellular processes including neuronal synaptic signaling.

## Introduction

WW domain, one of the smallest protein-protein interaction domains composed of only ∼35 amino acid residues, are widely distributed in many proteins with diverse and distinct functions (e.g. a totally of 95 WW domain in 53 proteins in the human proteome). WW domains organize molecular assemblies by binding to short, proline rich peptide motifs (∼4-6 residues) (Sudol and Hunter, 2000, Salah et al., 2012). Most of the WW domains in the human proteome belong to the type I (52 out of a total of 95). The type I WW domains bind to a consensus “PPxY” motif (where P is proline, x is any amino acid and Y is tyrosine, also known as PY-motif) with very modest affinities (with *Kd* in the range of a few to a few dozens of µM) (Chong et al., 2010, Kato et al., 2002, Kato et al., 2004, Aragon et al., 2011). The human proteome contains ∼1,500 “PPxY” in >1,000 proteins (Hu et al., 2004, Tapia et al., 2010). Therefore, the combination of potential WW domain/“PPxY” interactions are enormous. This immediately raises an issue on WW domain-mediated target binding specificities.

Taking the Hippo signaling pathway for an example, the pathway is organized by serval WW domain proteins (e.g. YAP, TAZ, KIBRA, and SAV1) and a number of PY-motif containing proteins such as LATS (LATS1 and LATS2), Angiomotins (AMOTs, including AMOT and AMOTL1/2), and PTPN14 (Pan, 2010, Sudol, 2010, Salah and Aqeilan, 2011, Yu and Guan, 2013) (Fig. 1A). The interactions between WW domains and PY-motifs in the Hippo pathway are very promiscuous, as nearly everyone of these WW containing proteins has been reported to interact with anyone of the PY-motif containing proteins. For example, YAP WW domains have been reported to bind to PY motifs from LATS, AMOTs, PTPN14, P73, SMAD1, etc. (Hao et al., 2008, Oka et al., 2008, Zhang et al., 2008, Chan et al., 2011, Wang et al., 2011, Zhao et al., 2011, Wang et al., 2012, Huang et al., 2013, Liu et al., 2013, Strano et al., 2001, Alarcon et al., 2009, Yi et al., 2013, Michaloglou et al., 2013). KIBRA WW domains have also been reported to bind to PY-motifs from LATS, AMOTs, and PTPN14 (Baumgartner et al., 2010, Genevet et al., 2010, Yu et al., 2010, Knight et al., 2018, Xiao et al., 2011, Hermann et al., 2018, Wang et al., 2014). The WW domains of SAV1 can bind to PY motifs of LATS (Tapon et al., 2002). In addition, PY motifs of LATS1, AMOT and PTPN14 also bind to WW domain-containing NEDD family E3 ligases (Kim and Jho, 2018, Nguyen and Kugler, 2018, Salah et al., 2012). However, it is not clear how such large array of WW/PY-motif interactions in the Hippo-related signaling pathway are inter-related during cell growth processes and whether all these reported interactions occur in living cells. Since whether YAP is in nuclei or in cytoplasm dictates the fate of cell growth and polarity (Sun and Irvine, 2016, Moya and Halder, 2018, Fulford et al., 2018, Yu et al., 2015a), it is envisaged that at least some of the WW/PY-motif interactions need to be specific.

**Figure 1.**
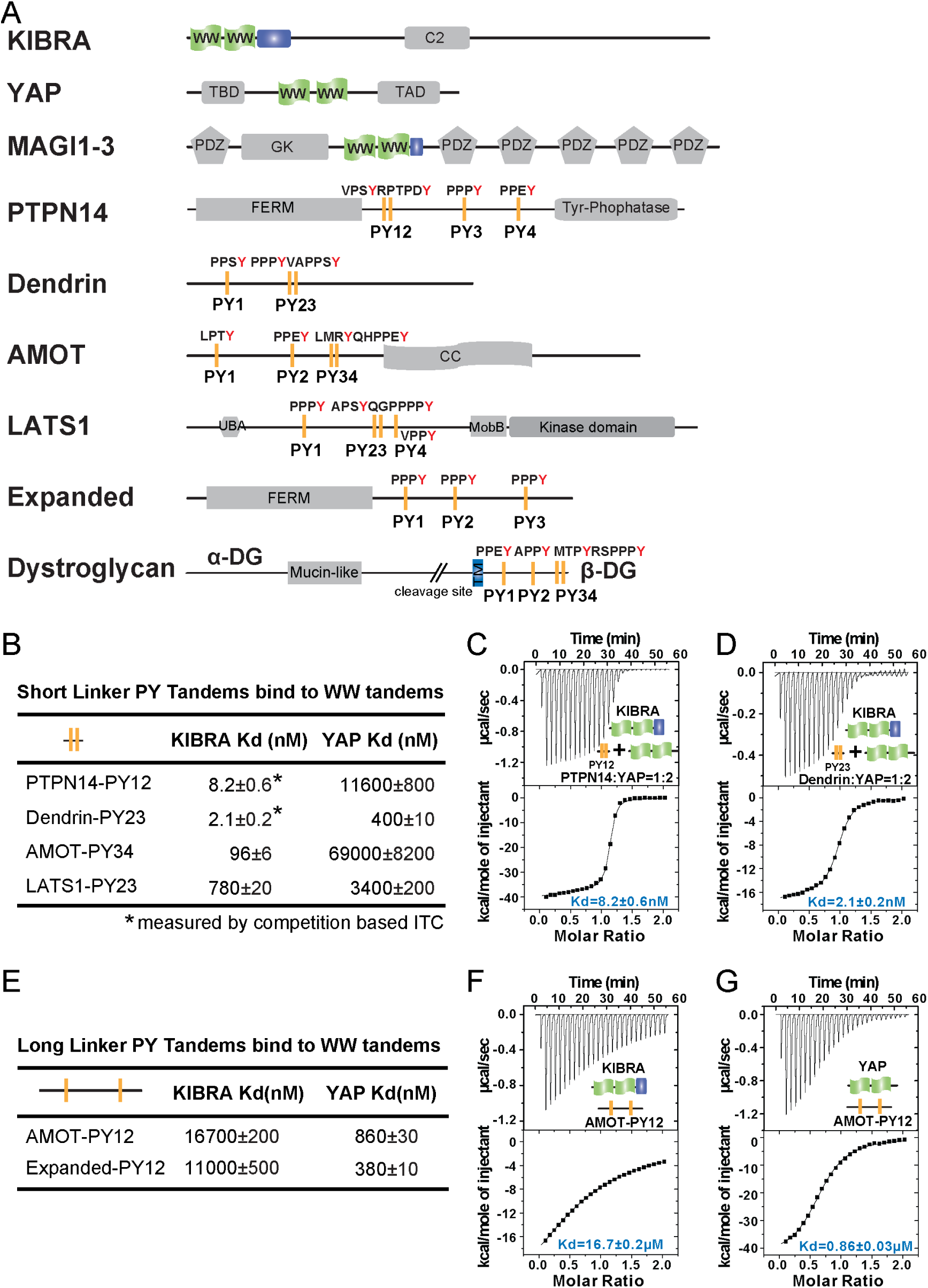
Super strong bindings of the KIBRA WW tandem to PY-motif containing proteins in cell growth control. (A) Schematic diagram showing the domain organization of selected WW-tandem containing proteins and multiple PY-motifs containing proteins in cell growth and polarity. (B) ITC-derived binding affinities of WW tandems from KIBRA and YAP in binding to PY motifs from PTPN14, Dendrin, AMOT and LATS1. (C-D) ITC-based measurements of the binding KIBRA WW12 to PTPN14 PY12 (C) and to Dendrin PY23 (D). Due to the very tight direct bindings between the WW tandem and each of the PY-motifs, we used a competition-based binding assay (i.e. by first saturated each PY-motifs with YAP WW12, which has a modest binding to the PY-motif) to obtain reliable dissociation constants. (E-G) ITC-based measurements of the bindings between WW tandems of KIBRA and YAP and PY motifs of AMOT and Expanded. Both PY motifs of AMOT and Expanded are separated by long and flexible linkers.

In cell growth and polarity regulations, YAP sits at the final converging point integrating both Hippo pathway-dependent and -independent upstream signals and regulating gene transcriptions via its WW tandem-mediated binding to PY-motif containing proteins including LATS, AMOTs and PTPN14 (Sun and Irvine, 2016, Moya and Halder, 2018, Fulford et al., 2018, Yu et al., 2015a). KIBRA also contains a WW tandem that binds to PY-motifs from the same set of proteins. Additionally, the cell polarity regulator, MAGIs (MAGI1-3), contains a WW domain tandem (Fig. 1A) that are likely capable of binding to various PY-motif proteins such as LATS, AMOTs and PTPN14 (Couzens et al., 2013, Wang et al., 2014, Bratt et al., 2005, Patrie, 2005). A systematic and quantitative study of the interactions between the three WW tandem proteins (YAP, KIBRA, and MAGIs) and many of the PY-motif containing proteins in cell growth control (e.g. those highlighted in Fig. 1A) will be vitally important to understand how these proteins function to orchestrate cell growth control.

In this study, we decoded the target binding mechanisms governing the WW tandem-mediated target interactions by focusing on proteins involved in cell growth and polarity regulations. To achieve this goal, we determined high reolustion structures of 7 representative WW tandem/target complex structures, and measured binding constants of numerous WW domain tandems or isolated WW domains binding to known or previously unknown targets. In sharp contrast to the common perception, we discovered that the WW tandem and PY-motif interactions are often extremely specific and strong, a finding that can rationalize numerous protein-protein interactions in cell growth and polarity. The rules underlying WW tandem/PY-motif interactions revealed in our study also allowed us to predict previously unidentified binders of WW domain proteins. The results presented in this study may serve as a portal for future studies of WW domain containing proteins in general.

## Results

### WW Tandem-mediated bindings to PY-motif proteins can be extremely specific and strong

To begin such systematic study, we first set out to compare bindings of the WW tandems of YAP and KIBRA with PY-motifs from LATS, AMOTs and PTPN14. We also included another PY-motif containing protein Dendrin in the study, serving as a reference for super strong binding involving WW domains (*Kd* ∼2.1 nM, Fig. 1D; also see our recent work by (Ji et al., 2019)). Similar to the two PY-motifs of Dendrin that are separated by only two residues, the first two PY-motifs of PTPN14 are also next to each other with only two residues in between, and the PY12 motif sequence is evolutionarily conserved (Fig. S1A). In contrast, PY3 and PY4 of PTPN14 are separated by 178 residues. We found that the KIBRA WW tandem also binds to PTPN14 with a super strong affinity (*Kd* ∼8.2 nM) (Fig. 1C). We were not able to perform quantitative binding assay between KIBRA WW tandem and PTPN14 PY34 as we could not obtain purified recombinant PY34. In a pull-down-based assay, interaction between the KIBRA WW tandem with PY34 was essentially undetectable when using PY12 as the control (data not shown). We further showed that KIBRA WW1 had a very weak binding to PTPN14 PY12 (*Kd* ∼32 μM) and KIBRA WW2 had no detectable binding to PTPN14 PY12 (Fig. S1B&C). Taken together, the above biochemical study revealed that the KIBRA WW tandem, via synergistic binding to the two PY-motifs immediately next to each other (i.e. PY12), can form a very tight complex with PTPN14.

The very tight bindings of the two closely spaced PY-motifs of PTPN14 (PY12) and Dendrin (PY23) with the KIBRA WW12 tandem (Fig. 1A&B) prompted us to analyze the PY-motif sequences of AMOTs and LATS carefully. We found that the last canonical PY-motif of AMOT (the “PPEY”-motif, named as PY3 in the literature) is preceded with a “LMRY”-motif, and these two motifs are only separated with two residues (Fig. 1A, and we define these two motifs as “PY34” from hereon). This sequence is conserved for AMOT and AMOTL1 throughout the evolution (Fig. S1A). We also found that the second canonical PY-motif of LATS1 (“PPPY”) is preceded with a “APSY”-motif, and the two motifs are separated by only three residues with the sequence of “^552^QGP^554^” (Fig. 1A, and these two motifs are defined as “PY23”). Again, the sequence containing the “APSY”- and “PPPY”-motifs and the gap residues in between are highly conserved in LATS1 in difference species (Fig. S1A). Interestingly, a Gly553Glu mutation of LATS1 was found in patients with renal cell carcinoma (Yu et al., 2015b). Somewhat unexpectedly, the KIBRA WW tandem binds to AMOT PY34 with a high affinity (*Kd* ∼96 nM), although the PY3 sequence severely deviates from the canonical WW domain binding PY-motif of “PPxY”. It is important to note that the KIBRA WW tandem binds to AMOT PY12, of which both PY1 and PY2 are canonical PY-motifs but are separated by 130 residues, only with a very weak affinity (*Kd* ∼16.7 μM) (Fig. 1E). We further found that the KIBRA WW tandem binds to LATS1 PY23 with a *Kd* of ∼0.78 μM (Fig. 1B), but much more weakly to the canonical PY1 motif (*Kd* ∼78 μM; Fig S1F). The above biochemical analysis revealed that, although one of the PY-motif deviates from the canonical WW domain binding PY-motif sequence, the two closely spaced PY-motifs in both AMOT and LATS1 can bind to the KIBRA WW tandem with high affinities.

The two WW domains of YAP are also arranged next to each other forming a WW domain tandem (Fig. 1A). Like the isolated WW1 in KIBRA, isolated WW1 or WW2 of YAP can bind to PY-motif sequences with *Kd* values of a few μM or a few tens of μM (Fig. S1). Very surprisingly, the YAP WW tandem binds to closely spaced PY-motifs of PTPN14, AMOT, and Dendrin with affinities of a few hundreds to a few thousands fold weaker than the KIBRA WW tandem does (Fig. 1B). These results revealed that WW domain tandem-mediated protein-protein interactions among proteins shown in Fig. 1A are extremely specific, a finding that is totally unexpected and will have immense implications in understanding the protein interactions and their functions in cell growth and cell polarity.

### Structures of the KIBRA WW tandem in complex with PY-motifs

We next pursued detailed structural studies trying to understand the mechanistic bases governing the vast affinity differences between the WW tandems of KIBRA and YAP in their bindings to the common set of PY-motif proteins shown in Fig. 1A. We solved the high-resolution crystal structures of KIBRA WW tandems in complex with PTPN14 PY12 (Fig. 2A), AMOT PY34 (Fig. 2B) and LATS1 PY23 (Fig. 2C; Table S1).

**Figure 2.**
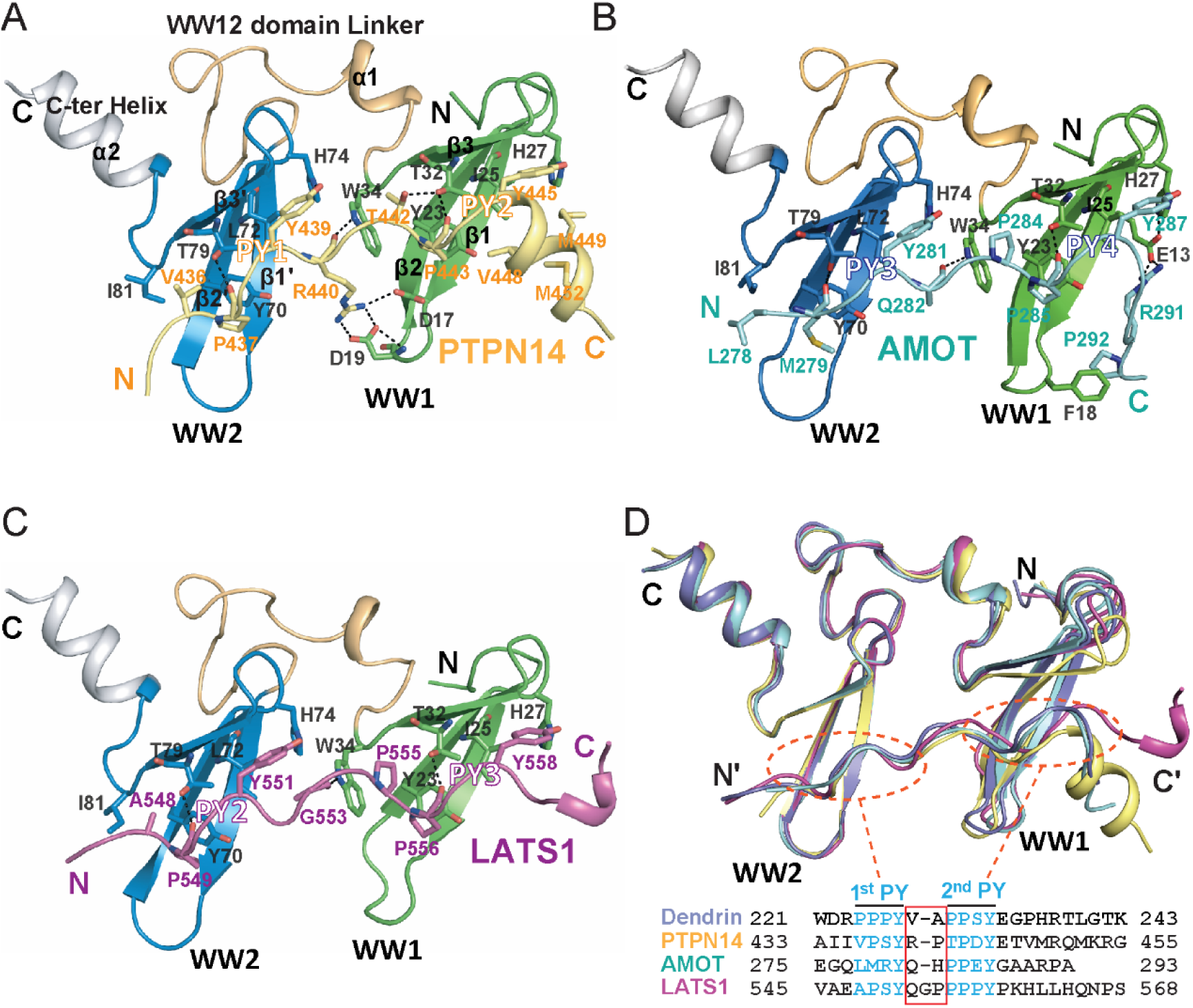
Structures of the KIBRA WW tandem in Complex with various closely spaced PY motifs. (A-C) Combined ribbon and stick model showing the structure and detailed interaction of the KIBRA WW tandem in complex with PY motifs from PTPN14 (A), AMOT (B), and LATS1 (C). (D) Superimposition of the structures of the KIBRA WW tandem in complex with PY motifs from PTPN14, AMOT, LATS1, and Dendrin. The amino acid sequences of the PY motifs are aligned at the bottom of the panel, and the red box highlights the 2-3 residue linking sequences of the PY motifs. The color-coding scheme of the structures is used throughout the manuscript.

The structures of the WW tandems in the three complexes are extremely similar, with the overall pairwise RMSD values in the range of 0.63-1.0 Å (Fig. 2D and S2A). Every WW domain forms a canonical β-sheet fold with three anti-parallel β-strands. The two WW domains align with each other in a head-to-tail manner, so that the two PY-motif binding pockets of the WW tandem are positioned right next to each other to accommodate the two PY-motifs in each of the three targets. The head-to-tail orientation of the WW tandem imposes a restriction that the target peptide can only bind to the WW tandem in an antiparallel manner (i.e. 1^st^ PY binds to WW2 and 2^nd^ PY binds to WW1; Fig. 2D). Though the direct contacts between the two WW domains in the tandem are minimal, the WW tandem forms a very stable structural supramodule by extensive interactions between the inter-domain linker (residues I35-L57) and an α-helix extension immediately following WW2 via hydrophobic (e.g. I35, P44, F47, A48, C50, L55, P56, L57 from the linker and W88 from α2), charge-charge and hydrogen bond interactions (e.g. the D53-R85-D83 network and Q80-Q87) (Fig. 3A). Perturbations of any of the inter-domain interactions (e.g. single point mutations such as I35D, F47A, L57D, and W88A, or deletion of the α2 helix) invariably weakened the binding of the WW tandem to its targets (Fig. 3B), indicating that formation of the WW12 supramodule is critical for high affinity bindings to the three targets studied here as well as to PY23 of Dendrin (Ji et al., 2019).

**Figure 3.**
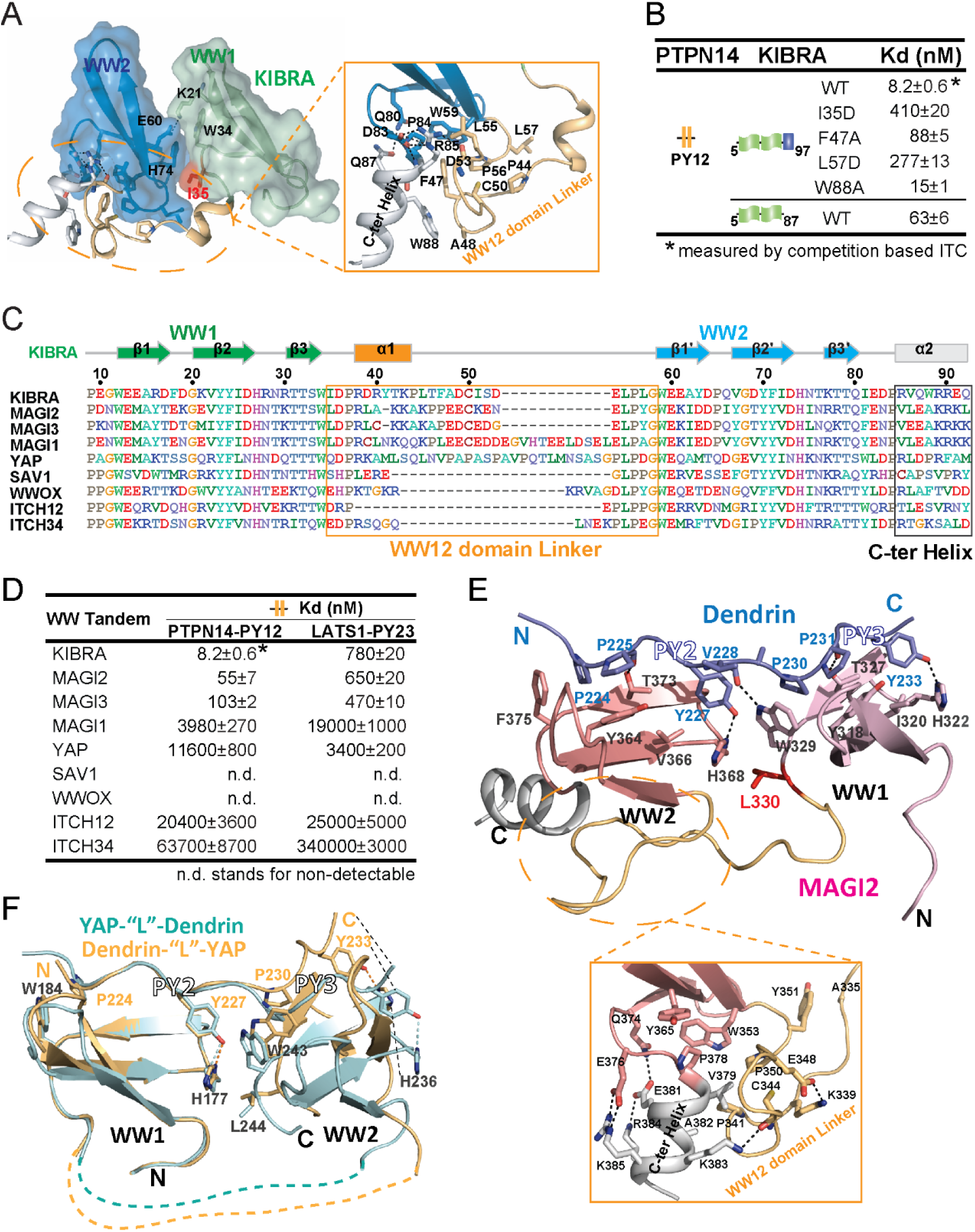
Structural basis governing the strong bindings of various WW tandems to their PY-motif containing targets. (A) Combined surface and ribbon-stick model showing the structural basis for the formation of the KIBRA WW tandem supramodule. The detailed interactions between the inter-domain linker and C-terminal α-helix are shown in an expanded insert at right. (B) ITC-based measurements comparing the binding affinities of the KIBRA WW tandem and its various mutants to PTPN14 PY12. (C) Amino acid sequence alignment of the WW tandems from KIBRA, MAGI1/2/3, YAP, SAV1, WWOX and ITCH-WW12/WW34, showing that the WW tandems of KIBRA, MAGI2, and MGAI3 are more similar to each other. (D) ITC-based measurements of the binding affinities of various WW tandems to PTPN14 PY12 and LATS1 PY23. (E) Combined ribbon and stick diagram showing the structure and detailed interaction of the MAGI2 WW tandem in complex with Dendrin PY23. The detailed interactions between the inter-domain linker and C-terminal α-helix are shown in an expanded box below the structure. (F) Ribbon and stick model showing the superimposition of two versions of the structures of YAP WW tandem in complex with Dendrin PY23. The structure colored in brown was obtained by fusing Dendrin-PY23 to the N-terminus of the YAP WW tandem, and the one colored in cyan was obtained by fusing Dendrin-PY23 to the C-terminus of the YAP WW tandem. The structures are superimposed by overlaying the WW1, showing different orientations of WW2 between the two structures.

The densities of all three ligands in the crystal structures, including the linker connecting the two PY-motifs, are well defined (Fig. S2B). Overall, the type I WW domain/“ΨPxY” sequence motif (where “Ψ” is an aliphatic or small polar amino acid with Pro being preferred) interaction mode is used for five of six WW/target interactions (AMOT’s 1^st^ PY motif has an unusual “LMRY” sequence, see below) in the three complexes (Fig. 2). In particular, the binding between WW1 and the second PY-motif of the three targets, each has a classical “ΨPxY”-motif (Fig. 2D), almost perfectly fits into the optimal type I WW/target binding (Fig. 2A-C) (Macias et al., 1996). The interactions between WW2 and the first PY-motif are more variable, as their sequences can significantly deviate from the optimal “PPxY”-motif (Fig. 2D). Correspondingly, KIBRA WW2 has an Ile (I81) instead of a highly conserved Trp at the end of the β3-strand (e.g. W34 in WW1; Fig. 2A-C). For example, the PY3-motif of AMOT has a sequence of “LMRY”, with the Leu and Met in the motif forming hydrophobic interaction with I81 and Y70 from WW2. This explains why KIBRA WW2 can accommodate non-“ΨPxY” motif sequences, as long as the two residues corresponding to “Ψ” and “P” are hydrophobic. At the same time, this non-canonical type I WW/target binding also explains that such interaction is weaker than the canonical type I WW/“PPxY”-motif, in which the two Pro residues in the motif are sandwiched by a Tyr in β2 and a Trp at the end of β3 (Y23 and W34 in WW1, Fig. 2B&C) forming a compact mini-hydrophobic core. Indeed, converting the first two residues in the non-ideal PY motif to the ideal “PPxY”-motif can significantly enhance the overall affinities of the interactions between the KIBRA WW tandem and two PY-motif-containing targets (see below). The fact that the type I WW domain can bind to a “ΦΦxY” sequence motif (where “Φ” can be any uncharged amino acids; e.g. “VPSY” and “TPDY” in PTPN14, “APSY” in LATS1, and “LMRY” in AMOT) significantly expands the target binding repertoire for WW domain-containing proteins.

In addition to the two PY-motifs, the residues in the linker and the C-terminal extension of the targets also contribute to their bindings to the KIBRA WW tandem. The linker residues R440 of PTPN14, Q282 of AMOT and G553 of LATS each use their carbonyl groups to form a hydrogen-bond with the sidechain of Trp34 (Fig. 2A-C). The sidechain of R440 of PTPN14 also forms charge-charge interactions with D17 and D19 in β12 loop of WW1 (Fig. 2A). The C-terminal extension of PTPN14 and that of AMOT form additional hydrophobic interactions with the WW1 domain (Fig. 2A&B). The residue R291 in AMOT C-terminal extension also interacts with E13 of WW1 domain through charge-charge interactions (Fig. 2B). All these interactions contribute to the bindings of PTPN14 and AMOT to KIBRA WW tandem.

### The WW tandems of MAGI2/3 and KIBRA resemble with each other in target bindings

Among the WW domain proteins, many contain WW domains connected in tandem (Fig. S3). We wondered whether there might exist type I WW tandems adopting similar high affinity target bindings as the KIBRA WW tandem does. Since the inter-domain linker and the α-helix following WW2 are critical for forming the supramodular structure and target bindings of the KIBRA WW tandem, we carefully searched for possible existences of these two elements in other type I WW tandems (Fig. 3C). By sequence alignment of WW tandems, including those from KIBRA, MAGIs, YAP, SAV1, WWOX, ITCH WW12 and ITCH WW34, we found that MAGI2/3 WW tandems have a striking similarity to the KIBRA WW tandem both in the linker (23 aa in MAGI2/3 vs 24 aa in KIBRA and with ∼60% sequence similarity) and the C-terminal helix regions (Fig. 3C and S4A). To our delight, ITC assays showed that the WW tandems of MAGI2 and MAGI3 indeed bind to PTPN14 PY12 with strong affinities (*Kd* values of ∼55 nM and ∼103 nM, respectively, Fig. 3D). It is noted that although MAGI1 WW tandem shares a similar C-terminal helix extension with the KIBRA WW tandem, it possesses a longer WW12 domain linker (Fig. 3C). Correspondingly, MAGI1 WW tandem was found to bind to PTPN14 PY12 with a modest affinity (*Kd* ∼4 μM; Fig. 3D), presumably due to spatial hindrance caused by the extra amino acids in the linker. We further found that MAGI2/3 WW tandems bind to Dendrin PY23 with extremely high affinities (*Kd* values of ∼2.3 and ∼3.6 nM, Fig. S4B), further supporting the conclusion that the MAGI2/3 and KIBRA WW tandems share a very high similarity in target bindings. Like KIBRA WW12, the MAGI2/3 WW tandem binds to LATS1 PY23 with modest affinities (Fig. 3D). Finally, the rest of type I WW tandems analyzed (i.e. those from YAP, SAV1, WWOX, ITCH WW12 and ITCH WW34; Fig. 3C) were found to bind to PTPN14 PY12 or LATS1 PY23 with very weak affinities or with no detectable binding (Fig. 3D). Taken together, the above biochemical and bioinformatics analyses strongly indicated that WW tandems (at least for the type I tandems investigated here) can have very strong affinities and exquisite specificities in recognizing their targets.

We next tried to determine the structures of MAGI2/3 WW tandems in complex with different PY tandems to gain further mechanistic insights into their target bindings. With extensive efforts, we were able to obtain high quality crystals of MAGI2 WW tandem in complex with Dendrin PY23 and determined the complex structure to the 1.65 Å resolution (Fig. 3E and Table S1). A structural comparison of the MAGI2 WW tandem in complex with Dendrin with those of the KIBRA WW tandem in complex with various ligands shown in Fig. 2 yielded RMSD values of 2.3-2.7Å for the Cα atoms, indicating that the structures of the KIBRA and MAGI2 WW tandems in these complexes are indeed very similar to each other (Fig. S4C). Notably, the extensive interactions between the inter-domain linker and the C-terminal helix stabilize the supramodular structure of the MAGI2 WW tandem, and the Dendrin PY23 binds to the MAGI2 WW tandem following almost the exactly same detailed interactions as those observed in the KIBRA WW12/Dendrin PY23 complex (Fig. 3E) (Ji et al., 2019).

### YAP WW tandem adopts a very different structure and target binding mode compared to the KIBRA WW tandem

We also tried to determine the structures of the YAP WW tandem in complex with various ligands listed in Fig. 1B to understand why YAP binds to these ligands with much lower affinities than does the KIBRA WW tandem. Despite extensive trials, we were not able to obtain any crystals when complexes were prepared by mixing the YAP WW tandem with the ligand peptides, presumably due to conformational flexibilities of the WW tandem (see below). We were, however, able to obtain high diffraction quality crystals when the Dendrin PY23 sequence was fused to either the N- or C-termini of the YAP WW tandem. We solved the structures of these two versions of the complex (Fig. 3F and S4D, Table S1). Although each WW domain engages the “PPxY” motif following the canonical type I WW/target bindings (Fig. S4D), the overall structure of the YAP WW tandem/Dendrin PY23 complex differs substantially from those of the KIBRA/MAGI2 WW tandems in complex with the same ligand (Fig. 3E *vs* 3F for example) in two distinct aspects. First, the inter-domain linker of the YAP WW tandem is unstructured and there is no C-terminal helix following YAP WW2. Thus, the two WW domains of YAP do not form a supramodule due to very limited inter-domain contact. This is supported by the two version of the complex structures solved, in which the inter-domain orientations are obviously different (Fig. 3F). Second, the Dendrin PY23 binds to the YAP WW tandem in a parallel orientation (i.e. PY2 binds to WW1 and PY3 binds to WW2), instead of the antiparallel binding orientation observed for the KIBRA/MAGI2 WW tandems. This is again likely attributed by the lack of inter-domain interactions between the two WW domains of YAP so that each domain can freely choose its more preferred PY-motif. The above structural analysis indicates that the YAP WW tandem would bind to the ligands containing two closely spaced PY motifs with only modest affinities due to the lack of conformational coupling of its two WW domains. The amino acid sequence linking the Yorkie WW tandem is totally different from that of YAP (Fig. S4E), further indicating that the linker sequence connecting the YAP WW tandem does not play active structural roles during the protein evolution. In contrast, the inter-domain linker sequences of the KIBRA and MAGI2/3 WW tandems are highly conserved during the evolution (Fig. S4A). Finally and as expected, the Yorkie WW tandem could only bind to Dendrin PY23 with a modest affinity (Fig. S4F).

### Origin of the exquisite target binding specificity of WW tandems

The similarities both in their structures and binding affinities to a set of common ligands indicate that the WW tandems of KIBRA and MAGI2/3 share a common binding mode (Fig. 4A, 4B and S4C). The extensive interactions between the inter-domain linker and the C-terminal helix couples the two WW domain into a structural supramodule in these WW tandems, thus providing a structural basis for highly synergistic bindings to target proteins. For example, the formation of the WW tandem supramodule allowed KIBRA WW12 to bind to PTPN14 with an ∼3900-fold higher affinity when compared to the isolated WW domains (Fig. 4D1 and S1B&C). In contrast, the coupling between WW1 and WW2 in YAP is minimal (Fig. 4C). Therefore, the target binding synergism between the two WW domains in the YAP tandem is also low. For example, the binding of YAP WW tandem to Dendrin PY23 is only a few dozen tighter than the isolated WW domains (Fig. 4D2 and S1B&D), and this relatively low synergism likely originates from the very closely spaced PY-motifs of Dendrin PY23. Consistent with this analysis, very little synergism could be observed in the bindings of the YAP WW tandem to two PY motifs separated with extended linking sequences such as AMOT PY12 (Fig. 4D3 and S1E).

**Figure 4.**
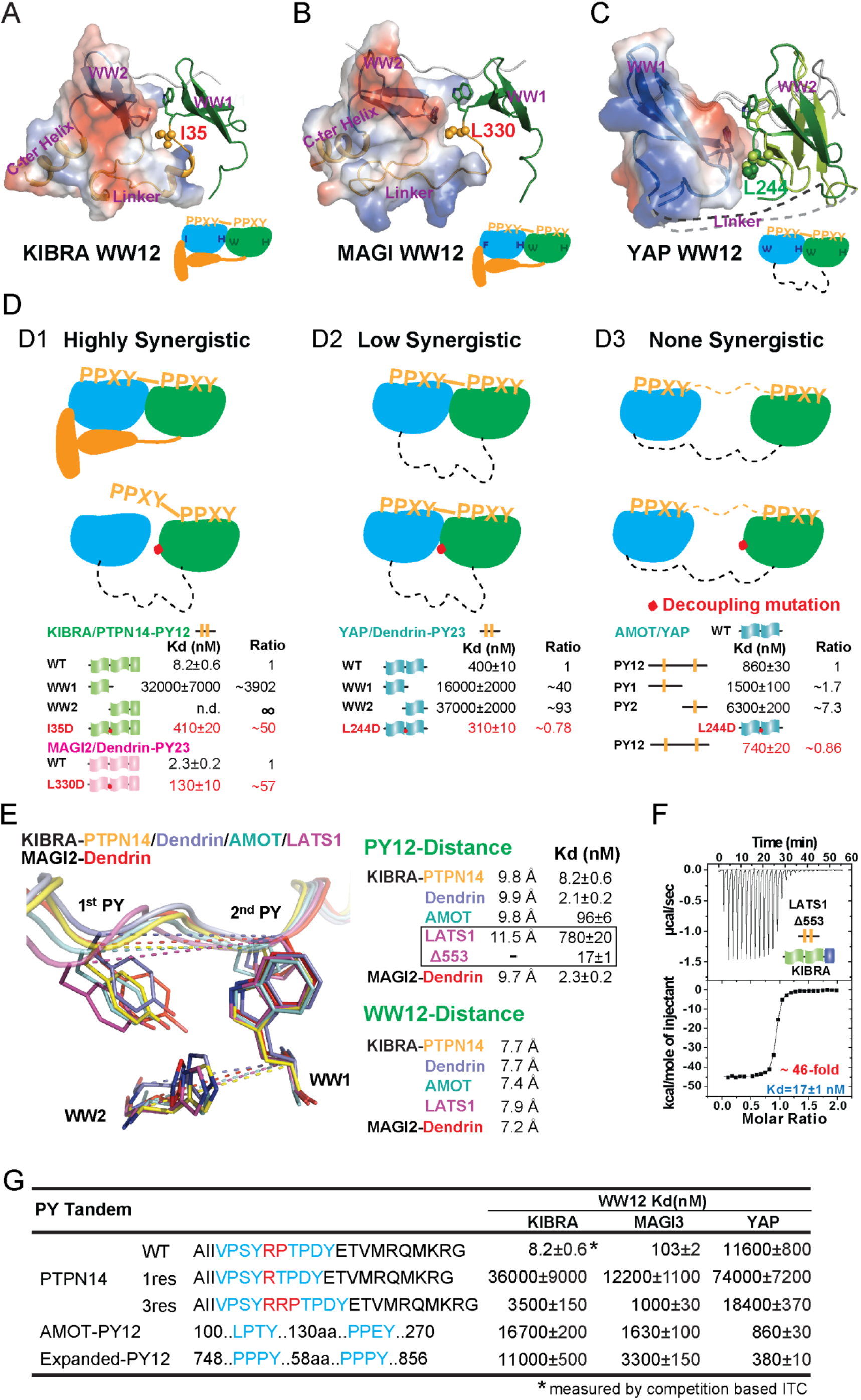
Binding modes of WW tandems to PY motifs. (A-C) Combined surface and ribbon diagram showing the domain coupling of the WW tandems of KIBRA (A), MAGI (B), and YAP (C), each in complex with Dendrin PY23. The cartoon in each panel is used to show the coupling mechanism of the WW domain tandems in the complexes. (D) Three distinct modes of WW tandem-mediated bindings to various PY motif targets. (D1), Highly synergistic bindings of tightly coupled WW domains in the tandem with two PY motifs separated by two and only two residues. In this mode, isolated WW domains have very weak binding to PY motifs as shown by the quantitative binding affinity data below the model cartoon. Perturbations of the inter-domain couplings (indicated by a red dot in the model) also severely impair the WW tandem/target bindings. (D2) Low synergistic binding of WW tandems with PY motif targets as illustrated by the two WW domains with no apparent direct coupling but binding to two PY motifs separated only by a very short linker. In this mode, isolated WW domains have very weak binding to PY motifs. Modest synergism can be brought to action by the two closely spaced PY motifs. Changes of WW domain surface outside the binding areas do not affect WW tandem/target bindings. (D3) None-synergistic bindings of WW tandems to PY motifs separated by long linkers. In this mode, bindings of the two WW domains to PY-targets are simply additive. (E) Ribbon-stick model showing the distances of the two PY motifs as well as two WW domains in the five complex structures. The inter-WW domain distances are indicated by the signature Trp at the end of β3-strand of WW1 and His in the β2-β3 hairpin of WW2. The distances between the PY motifs are measured Tyr in the first PY motif and the first Pro in the second PY motif. (F) ITC-based measurement showing that removal of one residue (Gly553) in the 3-residue linker (“QGP”) between LATS1 PY23 converted the LATS1 mutant into a super strong KIBRA WW tandem binder, presumably by shortening the inter-PY motif distance from 11.5 Å to ∼9.8 Å as shown in panel E. (G) ITC-derived binding affinities of various WW tandems to PTPN14 PY12 motif mutants with different inter-motif linker lengths. AMOT PY12 and Expanded PY12 are used as examples of PY motifs separated by longer linkers.

We directly tested the above WW domain coupling-induced target binding synergism model by experiments. In the WW tandem structures of KIBRA and MAGI2, a hydrophobic residue from WW1 (I35 in KIBRA and L330 in MAGI2) inserts into a hydrophobic pocket formed by WW2 domain and the inter-domain linker (Fig. 4A&B). Substitution of this hydrophobic residue with a negatively charged Asp should impair this hydrophobic interaction and thus exert a negative impact on the synergism of the two WW tandems. Indeed, the I35D substitution weakened KIBRA’s binding to PTPN14 PY12 by ∼50 fold, and the L330D mutation decreased MAGI2’s binding to Dendrin PY23 by ∼57 fold (Fig. 4D). L244 in YAP is the structurally equivalent residue of I35 in the KIBRA WW tandem (Fig. 4C). Totally consistent with the structural finding of very little coupling between the two WW domains in YAP, the L244D mutation had no impact on YAP’s binding to either Dendrin PY23 or AMOT PY12 (Fig. 4D). The L244D mutation data of YAP WW12 further support that the mild synergism of the binding between the YAP WW tandem and Dendrin PY23 originates from the two closely spaced PY motifs of Dendrin.

The strong structural couplings between the two WW domains of the KIBRA and MAGI WW tandems indicate that their two PY-motif-binding pockets are fixed both in distance and orientation. Indeed, the distances between Cα atoms of the Trp residue at the end of β3 of WW1 (one of the two defining Trp residues in WW domains) and of the His residue in β2-β3 hairpin of WW2 (which is absolutely required for engaging the Tyr residue in the “ΨPxY” motif for the type I WW domains) are in a very narrow range of 7.2-7.9 Å among the complex structures (Fig. 4E). Correspondingly, the distances between the two PY motifs of the ligands, measured between the Cα of Tyr in the 1^st^ PY and the Cα of Pro in 2^nd^ PY, are also fixed at ∼9.8 Å. The exception is the LATS1 PY23 motifs with the two Cα atoms separated by ∼11.5 Å (Fig. 4E). Notably, the PY-motifs in PTPN14, Dendrin and AMOT are all separated by only two residues, and all three ligands bind to the KIBRA and MAGI2/3 WW tandems with very high affinities. In contrast, the two PY-motifs of LATS1 are separated by three residues (“QGP”) and LATS1 binds to the two WW tandems with quite modest affinities (Fig. 3D). Strikingly, deleting the Gly residue (Gly553) in the three-residue linker of LATS1 PY23 converted the LATS1 mutant (“LATS1-Δ553”) into a very strong binder of the KIBRA WW tandem (*Kd* ∼17 nM; Fig. 4F), indicating that a two-residue linker between the two PY-motifs is optimal for specific and high affinity binding to the KIBRA and MAGI2/3 WW tandems. As expected, LATS1-Δ553 PY23 was found to bind to YAP WW12 with a similar affinity as the WT LATS1 PY23 does (data not shown). We further showed that shortening the linker of the PTPN14 PY12 to one residue (R, 1res) dramatically weakened its binding to both KIBRA and MAGI3 WW tandems (by as much as ∼4,400 fold; Fig. 4G), as two PY-motifs separated by only one residue would be shorter than the shortest distance between the two PY-motif binding pockets in the WW tandems (Fig. 4E). Similarly, lengthening the linker of the PTPN14 PY12 to three residues (RRP, 3res) also significantly weakened its bindings to the KIBRA and MAGI3 WW tandems (Fig. 4G). In contrast, lengthening the linker of the PTPN14 PY12 to three residues had negligible impact on its binding to the YAP WW tandem, as its conformational flexibility would allow the tandem to adjust the inter-domain distances and orientations.

In summary, we can divide the bindings between the WW domain tandems and two PY-motifs containing targets into three modes: 1), the highly synergistic and very strong binding mode, represented by the KIBRA and MAGI2/3 WW tandems, which requires the tight coupling of the two WW domains as well as two PY-motifs separated by two and only two residues (Fig. 4D1); 2), the low synergistic binding mode with modestly high affinities, represented by the YAP WW tandem, which does not require tight coupling between the two WW domains but with two PY-motifs separated by a short linker of a few residues (Fig. 4D2); and 3), those with no synergism and with weak binding affinities, in which the two WW domains do not couple with each other and the two PY motifs are separated by long linking sequences (Fig. 4D3).

### The deletion of Gly553 in LATS1 inhibits YAP phosphorylation in cells

According to the biochemical and structural analysis, deleting Gly553 from the LATS1 PY23 converted LATS1-Δ553 into a strong binder of KIBRA. We confirmed that KIBRA indeed binds to LATS1-Δ553 stronger than to WT LATS1 using a co-immunoprecipitation-based assay with the full-length proteins expressed in HEK293A cells (Fig. 5A, 5B). We further showed using purified recombinant proteins that KIBRA WW12 and YAP WW12 both formed complex with LATS1 PY1-4 on an analytical gel filtration column when the three proteins were mixed at a 1:1:1 ratio (Fig. S5A), fitting with the observation that the two WW tandems have comparable affinities in binding to LATS1 PY1-4 (Fig. S1F). In contrast, when YAP WW12, LATS1 PY1-4(Δ553) and KIBRA WW12 were mixed at a 1:1:1 ratio, essentially all KIBRA WW12 was found to be in complex with LATS1 PY1-4(Δ553) and YAP WW12 was found to be competed off and existing in the free form (Fig. S5B). Interestingly, when Gly553 of LATS1 was substituted with Glu, the mutant LATS1 showed very weak binding to KIBRA (Fig. S5D), and all KIBRA WW12 was found to be in the free form and most of YAP WW12 formed complex with the LATS1 mutant when KIBRA WW12,YAP WW12 and LATS1 PY1-4(G553E) were mixed at a 1:1:1 ratio (Fig. S5C). The weakened binding of the LATS1-G553E mutant to KIBRA WW12 is likely due to more rigid backbone conformation of Glu over the Glycine residue in the WT protein, such that the defined distince between the two PY-motif binding pockets of KIBRA WW12 can no longer accommodate the LATS1 PY23 mutant.

**Figure 5.**
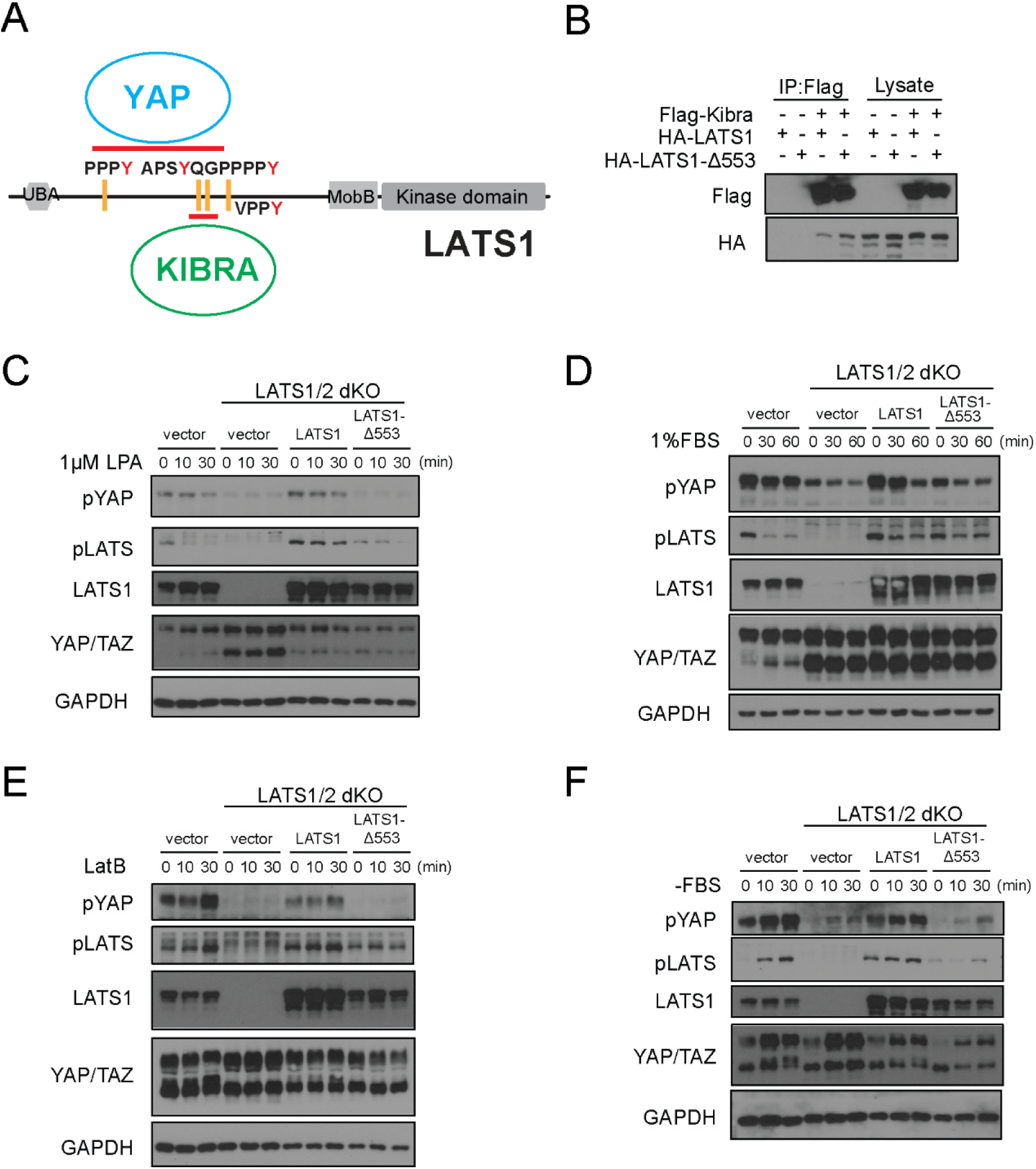
Deletion of Gly553 in LATS1 inhibits YAP phosphorylation in cells. (A) Schematic diagram showing the interaction of LATS with KIBRA or YAP. (B) LATS1-ll553, compared to WT LATS1, has an enhanced binding to KIBRA in HEK293A cells. (C, D) Comparison of YAP and LATS1 phosphorylations of wild type HEK293A cells, LATS1/2 dKO cells, LATS1/2 dKO cells stably expressing LATS1 or LATS1-ll553. Cell were serum-starved for 4h and then treated with 1 μM LPA (C) or 1%FBS (D) for indicated times. (E, F) Comparison of YAP and LATS1 phosphorylations of wild type HEK293A cells, LATS1/2 dKO cells, LATS1/2 dKO cells stably expressing LATS1 or LATS1-ll553 treated with 0.2 mg/ml LatB (E) or serum starvation (F) for indicated times.

We reasoned that the dramtically enhanced binding between KIBRA and LATS1-Δ553 would shift the LATS1 mutant from the YAP/LATS complex to the KIBRA/LATS complex and thus diminish the LATS-induced YAP phosphorylation when cells are stimulated with the Hippo pathway activation signals. To test this hypothesis, we used the LATS1/2 dKO HEK293A cells that we characterized earlier (Plouffe et al., 2016). We generated stable cell pools by re-expressing WT LATS1 or LATS1-Δ553 in the the LATS1/2 dKO cells to test YAP and LATS1 phosphorylations upon various Hippo pathway stimulation signals. Upon treatment with lysophosphatidic acid (1 µM LPA), serum stimulation (1% FBS), latrunculin B (LatB, 0.2 mg/ml), or serum starvation (−FBS), which are well-known signals for the Hippo pathway (Yu et al., 2012, Meng et al., 2015), cells expressing LATS1-Δ553 had diminished YAP and LATS1 phosphorylations when compared with cells expressing WT LATS1 (Fig. 5B-D). These data suggest that the exquisite specificities of the WW domain tandem-mediated target bindings are critical for proper Hippo pathway signaling.

### Further specificity determinants of PY-motifs for highly synergistic bindings to WW tandems

Several additional sequence features besides the two-residue linker connecting the two PY-motifs contribute to the very high affinity bindings to the tightly coupled WW tandems. First, residues in the two-residue linker can enhance binding. For example, R440 of PTPN14 PY12 forms salt bridges with D17 and D19 in the WW1 of the KIBRA tandem (Fig. 2A). Substitution of R440 with Ala weakened PTPN14’s binding to KIBRA by 17-fold (Fig. 6A&B). Second, the C-terminal extension of PTPN14 PY12 tandem enhances its binding to KIBRA WW tandem by forming hydrophobic interactions with its WW1 domain (Fig. 6A). Truncating this C-terminal extension weakened PTPN14 PY12’s binding to KIBRA WW tandem by 22-fold (“Del-C” in Fig. 6B). Such C-terminal extension-enhanced binding also occurs for the binding of PTPN14 PY12 to MAGI3 WW tandem (Fig. 6B). It is also noted that LATS1 PY23 does not have the C-terminal extension, and it has weaker affinities in binding to KIBRA and MAGI WW tandems (Fig. S4B). Third, converting the non-canonical PY-motifs into canonical ones (i.e. “PPxY”) further enhanced PTPN14 PY12’s binding to both KIBRA and MAGI3 WW tandems by about 10- and 25-fold, respectively (“2Pro” in Fig. 6B). Further replacements of the residue in the “x” position of the canonical PY-motifs with Pro enhanced PTPN14 PY12’s binding to KIBRA and MAGI3 WW tandems by a modest ∼2-4 folds (“4Pro” in Fig. 6B), possibly due to further backbone conformation rigidification. Finally, although not tested here, one may be able to further enhance the ligand binding to the KIBRA and MAGI WW tandems by utilizing a small unoccupied hydrophobic surface of WW2 to bind to residue(s) N-terminal to the first PY-motif (highlighted with a dashed oval in Fig. 6A)(Qi et al., 2014, Liu et al., 2016).

**Figure 6.**
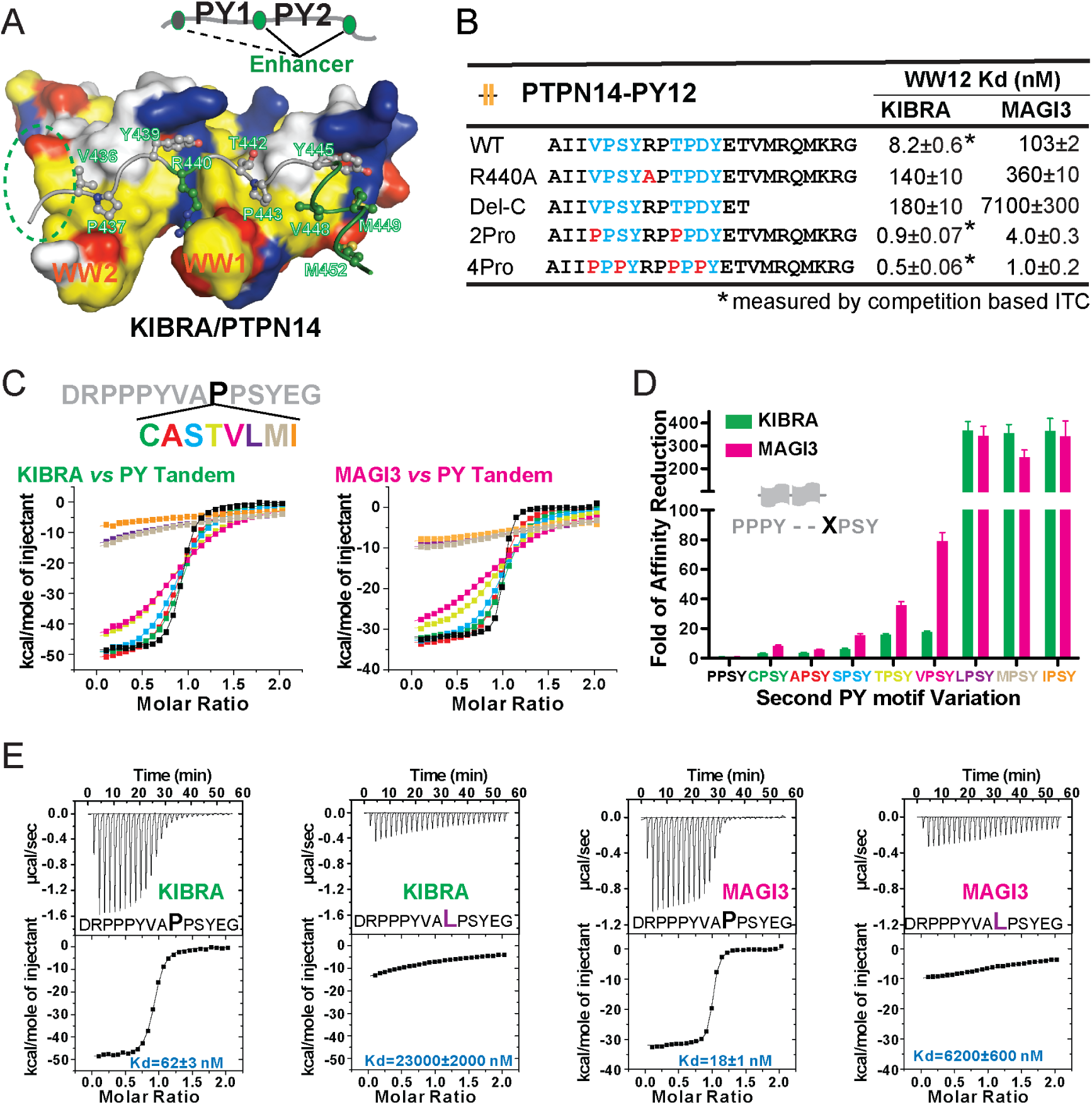
Specificity determinants of closely spaced PY motif to the WW tandems of KIBRA and MAGI2/3. (A) Surface combined with ribbon-stick model showing the binding between the KIBRA WW tandem and PTPN14 PY12. The figure is to show that, in addition to the WW/PY motif interactions, the inter-PY motif linker and the extension sequence following PY2 (as indicated by a cartoon) also play critical roles for the super strong bindings between the WW tandem and the PY motifs. In the drawing, the hydrophobic, negatively charge, positively charged, and polar residues are colored in yellow, red, blue and white, respectively, in the surface model. (B) ITC-derived binding affinities of the KIBRA and MAGI3 WW tandem to various PTPN14 PY12 mutants. For bindings with nM or sub-nM *Kd* values, competition-based ITC assays were used as described in Fig. 1C&D. (C) Mutational analysis of the contributions of different amino acids corresponding to the first Pro in the second PY motif of Dendrin PY23 in binding to KIBRA and MAGI3 WW tandems. The figure shows the superposition plot of ITC-based binding curves of binding reactions performed under the same experimental condition. (D) Comparison of relative binding affinity difference of various PY motif mutants tested in Panel C. In this comparison, the bindings of the KIBRA and MAGI3 WW tandems to the wild type PY motifs with the optimal “PPSY” sequence are set at the base value of “1”. The fold of affinity reductions of each mutant are plotted. (E) The representative ITC curves of showing the binding of the KIBRA/MAGI3 WW tandems to the WT PY motif peptide or to one of the representative mutants.

The finding that canonical WW domains can bind to non-canonical PY-motif sequences significantly expands possible WW domain/target binding space. It is thus important to define what residues are allowed in the first position of the “ΨPxY” motif in addition to Pro for binding to canonical WW domains. We modified the first Pro of the second PY in the peptide “DRPPPYVA**<u>P</u>**<u>PSY</u>EG” (the bold face “P” in the Dendrin PY23 peptide without the C-terminal extension) with Leu, Ala, Val, Cys, Ser, Ile, Thr, and Met, and measured the bindings of these peptides to the KIBRA and MAGI3 WW tandems by ITC (Fig. 6C). As expected, all substitutions weakened the peptide binding to the tandems (Fig. 6C-E, S6). However, substitutions of the Pro residue with Cys, Ala, Ser, Thr, and Val lead to only mild weakening of the binding, whereas substitutions with bulky hydrophobic Leu, Met and Ile essentially disrupted the binding. The above results can be rationalized by the structure of the KIBRA WW tandem in complex with PTPN14 PY12 showing that the pocket accommodating the residue (T442 in Fig. 6A) is quite small.

Our structural studies of the five WW tandem/target complexes also revealed that the sequence requirement for the first PY motif is relatively loose. Though the “PPxY” sequence is optimal (Fig. 6B), the two Pro residues can be other aliphatic residues or even uncharged polar amino acids (Fig. 2). This can be explained by the relatively shallow and open hydrophobic pocket that the WW2 domain uses to accommodate the first two residues from the first PY motif (e.g. V436 and P437 in Fig. 6A). Taken all above results together, we can deduce a consensus sequence motif for specific and strong bindings to the KIBRA/MAGI2/3 WW-tandems as: “ΦΦxYxxΨPxY”, where Φ is aliphatic or small polar amino acids and Ψ can be Pro, Cys, Ala, Ser, Thr or Val (Fig. 7A). The first PY-motif of such sequence binds to WW2 and the second PY-motif binds to WW1 of the WW tandems (Fig. 7B).

**Figure 7.**
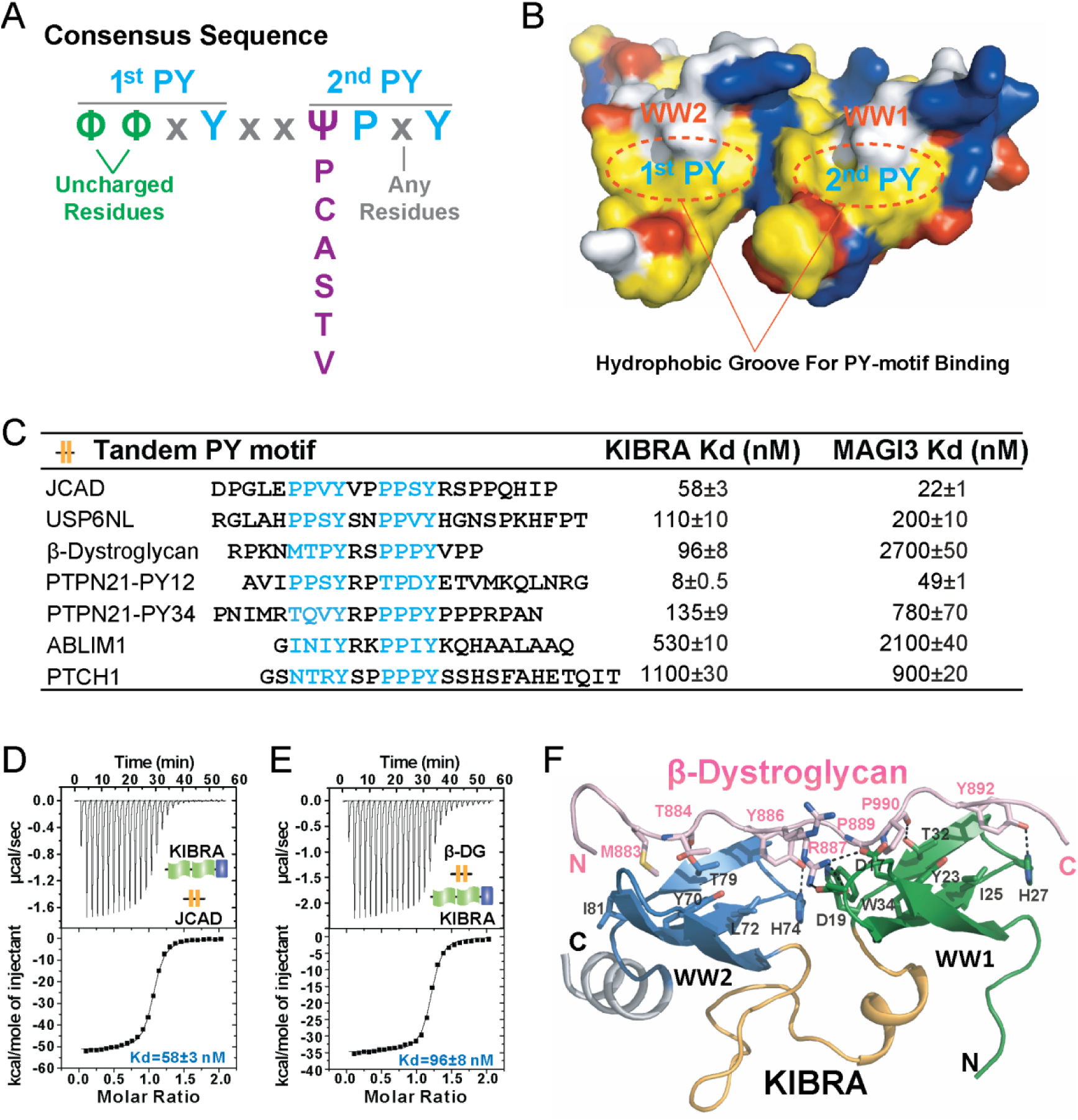
A consensus sequence of closely-spaced PY motifs capable of binding to KIBRA and MAGI2/3 WW tandems with high affinities. (A) The consensus sequence pattern of two PY motif-containing sequences that can synergistically bind to tightly couple WW tandems such as those from KIBRA and MAGI2/3. (B) Surface diagram showing two hydrophobic PY motif binding pockets of WW tandem that are precisely spaced due to the tight inter-domain coupling. The color coding is the same as in Fig. 5A. (C) ITC-based measurements of the binding affinities of selected targets with PY motifs fitting the consensus sequence indicated in panel A to WW tandems from KIBRA and MAGI3. (D&E) Representative ITC binding curves of the bindings described in panel C, showing the bindings of KIBRA WW tandem to PY motifs JCAD (D) and β-Dystroglycan (E). (F) Ribbon-stick model showing the structure and detailed interaction of the KIBRA WW tandem in complex with the PY motifs from β-Dystroglycan.

### Identification of new binders of the KIBRA and MAGI2/3 WW tandems

We searched for the proteins in the UniProtKB/Swiss-Prot database (UniProt-Consortium, 2019) and identified 132 human proteins containing the “ΦΦxYxxΨPxY” motifs (Table S2). Besides Dendrin, only two proteins, junctional protein associated with coronary artery disease (JCAD) and USP6 N-terminal-like protein (USP6NL), contain two canonical “PPxY” motifs separated by exactly two residues (Fig. 7C). PTPN21 contains two consensus sequences with its PY12 highly similar to PTPN14 PY12 and PY34 being unique to PTPN21 itself (Fig. 7C).

We selected 7 of such consensus sequences from 6 proteins (JCAD, USP6NL, β-Dystroglycan, PTPN21, ABLIM-1, and PTCH1) to measure their bindings to the KIBRA and MAGI3 WW tandems by ITC (Fig. 7C and S7). As expected, both JCAD and USP6NL bind strongly to the KIBRA WW tandem (Fig. 7C&D). Both PY12 and PY34 of PTPN21 show strong bindings to the KIBRA WW tandem. PTPN21 PY12 and PTPN14-PY12 bind to KIBRA WW tandem with essentially the same affinity (*Kd* ∼8 nM), as the sequences of these two peptides are very similar. Interestingly, β-Dystroglycan, a protein known to play critical roles in Hippo signaling-related cell adhesion and cell growth (Morikawa et al., 2017, Gawor and Proszynski, 2018), can also bind to KIBRA WW tandem with high affinity (*Kd* ∼96 nM, Fig. 7E). We solved the crystal structure of the KIBRA WW tandem in complex with the β-Dystroglycan PY34 peptide. The complex structure confirms that the peptide binds to the KIBRA WW tandem following the mode exactly as we have predicted (Fig. 7F). Therefore, β-Dystroglycan may be a key regulator in cell growth and neuronal synaptic functions via specifically binding to KIBRA.

## Discussion

In this study, we discovered that WW tandems, but not individual WW domains, can bind to target proteins with extremely high affinity (*Kd* in the range of low nM) and specificity. Such strong bindings are realized by both highly synergistic conformational coupling of two WW domains connected in tandem and two PY-motifs separated by a two and only two amino acid residues. If only using either one of the above two elements, the bindings become to be weakly synergistic and with modest affinities with *Kd* of a few hundreds of nM to a few µM. When two uncoupled WW domains binding to two PY motifs separated by more than two residues, their bindings are weak with *Kd* of a few to a few tens µM or even weaker due to very little or no synergism of the bindings between the two domains (Fig.4D). Weak targeting binding synergisms resulted from week couplings of WW domains connected in tandem have also been reported in several other cases in the literature (Aragon et al., 2011, Verma et al., 2018, Fedoroff et al., 2004, Liu et al., 2016, Qi et al., 2014). Additionally, it is well known that isolated WW domains bind to their targets with low affinities (*Kd* in the range of a few tens to a few hundreds of µM) (Sudol and Hunter, 2000, Salah et al., 2012). Therefore, WW domains can bind to their target proteins with a very broad affinity range with *Kd* values range from a few nM to hundreds of µM or weaker. Presumably, those highly affinity WW tandem-mediated target interactions should play specific functions. For example, we recently showed that the tight binding between KIBRA and Dendrin (*Kd* ∼2.1 nM) is critical for KIBRA (and its associated AMPA receptors) localization in neuronal synapses and for synaptic plasticity (Ji et al., 2019). We have also demonstrated that bindings of short PY motif containing peptides to WW tandems can be readily enhanced to sub-nanomolar affinities (Fig. 6B), and these super-strong WW tandem binding peptides may be used as effective tools for studying the role of specific interactions between WW tandem proteins and target proteins (see (Ji et al., 2019) for an example). As for the very weak WW domain/target interactions, cautions are required in interpreting potential functional implications of such bindings.

As for the WW domain-mediated protein-protein interactions in cell growth and polarity, the results presented in the current study will be valuable to re-evaluate many of reported protein-protein interaction and for guiding experimental design in the future. For example, the bindings of the KIBRA and MAGI tandems to PTPN14 (and PTPN21) are so much stronger than the binding of YAP to PTPN14 (∼1,400-fold difference; Fig. 1B). Therefore, it is almost for sure that PTPN14 should first bind to KIBRA and MAGI instead of to YAP when PTPN14 is not in excess. The binding between PTPN14 to YAP can occur if PTPN14 is in excess of total amount of KIBRA and MAGI together. Alternatively, PTPN14 may use its PY34 to bind to the YAP WW tandem via rather weaker bindings (Wang et al., 2012, Huang et al., 2013, Liu et al., 2013, Michaloglou et al., 2013). Similarly, the binding of the KIBRA tandem (or MAGI2/3 WW tandems) to LATS is ∼4.5-fold stronger than YAP tandem to LATS (Fig. 3D and Fig. S1F), therefore KIBRA can effectively modulate the concentration of the LATS/YAP complex formation by competing with YAP for binding to LATS. This is supported by the finding that further enhancing KIBRA/LATS1 binding by deleting Gly553 in the LATS1 PY23 linker can effectively block LATS1-mediated YAP phosphorylation, presumably due to effective sequestration of LATS1-Δ553 by KIBRA (Fig. 5). It should be noted that there are multiple strong KIBRA WW tandem binders such as PTPN14, PTPN21, Dendrin, AMOT, β-Dystroglycan, and JCAD (Fig. 1B and 7C), these molecules can tune the availability of free KIBRA in competing with YAP for binding to LATS. One may view that proteins like KIBRA and MAGIs function as an upstream nexus that can sense various cell growth and polarity cues and then determine whether and how much the LATS/YAP complex can form (i.e. determining the level of YAP phosphorylation by LATS) in living tissues. Conversely, certain PY motif containing proteins such as AMOTs can effectively compete with LATS for binding to YAP (Fig. 1E) and thereby regulate the LATS/YAP complex formation. It is noted that most of studies in elucidating molecular roles of WW domain proteins and their PY motif-containing targets involved in cell growth and cell polarity in the literature used protein overexpression or removal approaches. Such methods unavoidably would alter the WW domain-mediated protein-protein interaction network, which is very sensitive to the concentrations of each protein based on the quantitatively binding affinity data presented in this work. Therefore, we advocate that, when studying proteins in cell growth and polarity, cares should be taken to consider the quantitative binding affinities and endogenous concentrations of these proteins.

## Methods

### Constructs, Protein Expression and Purification

The coding sequences of desired constructs were PCR amplified from mouse brain cDNA libraries. For crystallization, Dendrin (222-241) was covalently linked at N- or C-terminal of YAP (156-247) by a two-step PCR-based method (referred to as Dendrin- “L”-YAP and YAP-“L”-Dendrin). All site mutations were created by PCR and confirmed by DNA sequencing. All constructs used for protein expression were individually cloned into a pET vector and recombinant proteins were expressed in Codon-plus BL21 (DE3) *Escherichia coli* cells with induction by 0.3 mM IPTG at 16 °C overnight. All recombinant proteins were purified using Ni^2+^-nitrilotriacetic acid agarose (Ni-NTA) column followed by size-exclusion chromatography (Superdex 200 column from GE Healthcare) in a final buffer containing 50 mM Tris-HCl (pH 7.8), 100 mM NaCl, 1 mM DTT and 1 mM EDTA. The selenomethionine-labeled proteins were expressed by methionine auxotroph *E. coli* B834 cells in LeMaster media and was purified following the same protocol used for the wild-type proteins. All tags of recombinant proteins were removed before crystallization, except for the His6-tagged YAP-“L”-Dendrin.

### Isothermal Titration Calorimetry Assay

Isothermal Titration Calorimetry (ITC) experiments were carried out on a VP-ITC calorimeter (MicroCal) at 25°C. Titration buffer contained 50 mM Tris-HCl (pH 7.8), 100 mM NaCl, 1 mM DTT and 1 mM EDTA. For a typical experiment, each titration point was performed by injecting a 10 μL aliquot of protein sample (50-300 μM) into the cell containing another reactant (5-30 μM) at a time interval of 120s to ensure that the titration peak returned to the baseline. For the competition experiments, proteins in the syringe were titrated to a mixture with a 2-fold molar concentration excess of a competitor over the reactants in the cell. The titration data were analyzed with Origin7.0 (MicroCal) using a one-site binding or competitive binding model.

### Crystallization and Data Collection

All crystals were obtained by hanging drop vapor diffusion methods at 4 °C or 16 °C within 3-5 days. Crystals of wild-type and Se-Met-substituted KIBRA/LATS1 were grown in solution containing 20-30% w/v PEG 4000 and 100 mM Tris (pH 8.0); crystals of KIBRA/AMOT were grown in solution containing 20-25% w/v pentaerythritol propoxylate 629 (17/8 PO/OH), 50 mM MgCl2 and 100 mM Tris (pH 8.5); crystals of KIBRA/PTPN14 were grown in solution containing 700 mM magnesium formate and 100 mM Bis-Tris Propane (pH 7.0); crystals of KIBRA/β-Dystroglycan were grown in solution containing 3.0-4.0 M NaCl and 100 mM Bis-Tris (pH 6.5); crystals of MAGI2/Dendrin were grown in solution containing 2.0-2.5 M ammonium sulfate and 100 mM sodium acetate (pH 4.6); crystals of Dendrin-“L”-YAP were grown in solution containing 28-35% w/v pentaerythritol propoxylate 426 (5/4 PO/OH) and 100 mM HEPES (pH 7.5); and crystals of His6-tagged YAP-“L”-Dendrin were grown in solution containing 25-30% w/v PEG400, 200 mM CaCl2 and 100 mM HEPS (pH 7.5). Before diffraction experiments, crystals were soaked in the original crystallization solutions containing an additional 10-25% glycerol for cryoprotection. All diffraction data were collected at the Shanghai Synchrotron Radiation Facility BL17U1 or BL19U1. Data were processed and scaled by HEL2000 or HKL3000.

### Structure Determination and Refinement

For the KIBRA/LATS1 complex, the program HKL2MAP (Pape and Schneider, 2004) yielded two Se sites in one asymmetric unit. The initial SAD phases were calculated using PHENIX (Adams et al., 2010). Other structures were determined by molecular replacement using PHASER (McCoy et al., 2007). The initial phases of KIBRA/ligands complex were solved using the WW tandem part from the structure of KIBRA/LATS1 complex as the searching model. The initial phase of MAGI2/Dendrin complex was solved using the structures of MAGI1-WW1 (PDB: 2YSD) and MAGI1-WW2 (PDB: 2YSE) as the searching models. The initial phases of His6-tagged YAP-“L”-Dendrin was solved using the structures of YAP-WW1 (PDB: 4REX) and YAP-WW2 (PDB: 2L4J) as the searching models. The initial phases of Dendrin-“L”-YAP was solved using the structures of YAP-WW1/2 from the structure of YAP-“L”-Dendrin. PY tandems were manually built according to the *F*obs-*F*calc difference Fourier maps in COOT (Emsley et al., 2010). Further manual model adjustment and refinement were completed iteratively using COOT (Emsley et al., 2010) and PHENIX (Adams et al., 2010) based on the 2*F*obs-*F*calc and *F*obs-*F*calc difference Fourier maps. The final models were further validated by MolProbity (Williams et al., 2018). Detailed data collection and refinement statistics are summarized in Table S1. All structure figures were prepared using PyMOL (http://pymol.sourceforge.net/).

### Analytical gel filtration chromatography

Analytical gel filtration chromatography was carried out on an AKTA FPLC system (GE Healthcare). Protein samples (each protein at a concentration of 40 μM) were loaded to a Superose 12 10/300 GL column (GE Healthcare) pre-equilibrated with assay buffer (same with ITC).

### Cell culture, transfection and retroviral infection

All cell lines were maintained in DMEM (GIBCO) containing 10% fetal bovine serum (FBS, GIBCO), 100 U/mL penicillin and 100 μg/mL streptomycin and with 5% CO2 at 37 C. To stimulate the Hippo pathway, low confluence cells (1.0 × 10^5^ cells per well) in 12-well plates were treated with serum stimulation, serum starvation, LPA (1 μM), and LatB (0.2 mg/ml).

The human LATS1-Δ553 plasmid was cloned by PCR and confirmed by DNA sequencing. Cells were seeded in 6-well plates for 24 hours, and then were transfected with plasmids using PolyJet Reagent (Signagen Laboratories) according to manufacturer’s instruction. To generate HEK293A cells stably expressing WT LATS1 or LATS1-Δ553, stable HEK293A cells with LATS1/2 dKO(Plouffe et al., 2016) were individually infected with retrovirus packed with plasmid expressing empty vector (pQCXIH), LATS1, or LATS1-△553. Fourty-eight hours after infection, cells were selected with 250 μg/mL hygromycin (Roche) in the culture medium.

### Immunoblotting and Immunoprecipitation

Immunoblotting was performed with the standard methods as described (Yu et al., 2012, Meng et al., 2015). Antibodies for pYAP(S127)(#4911), LATS1(#3477), pLATS(#8654S), HA-HRP(#2999) were from Cell Signaling Technology; Anti-Flag antibody (#F1804) was from Sigma-Aldrich; for YAP/TAZ(#sc-101199) was from Santa Cruz Biotechnology.

For immunoprecipitaion, cells were harvested at 48 hrs after transfection with lysis buffer containing 50 mM Tris (pH 7.5), 150 mM NaCl, 1 mM EDTA, 1% NP-40, 10 mM pyrophosphate, 10 mM glycerophosphate, 50 mM NaF, 1.5 mM Na3VO4, protease inhibitor cocktail (Roche), and 1 mM PMSF. After centrifuging at 13,300×g for 15 min at 4℃, supernatants were collected for immunoprecipitation. Anti-flag antibody was added to the supernatants and the mixtures were incubated overnight at 4℃. Then Pierce™ Protein A/G Magnetic Beads were added to the mixtures and further incubated for 2 hrs at 4℃. Immunoprecipitate proteins were eluted with SDS-PAGE sample buffer, reolved by SDS-PAGE and probed by the anti-HA antibody.

### Data Resources

The atomic coordinates of the WW tandem and target complex structures have been deposited to the Protein Data Bank under the accession codes of: 6J68 (KIBRA/LATS1), 6JJW (KIBRA/PTPN14), 6JJX (KBIRA/AMOT), 6JJY (KIBRA/β-DG), 6JJZ (MAGI2/Dendrin), 6JK0 (YAP-Linker-Dendrin), and 6JK1 (Dendrin-Linker-YAP).

## Acknowledgments

We thank the BL19U1 beamline at National Facility for Protein Science Shanghai (NFPS) and BL17U1 beamline at Shanghai Synchrotron Radiation Facility (SSRF) for X-ray beam time. This work was supported by a National Key R&D Program grant of China (2016YFA0501903) from the Minister of Science and Technology of China, grants from RGC of Hong Kong (AoE-M09-12 and C6004-17G), a grant from Asia Foundation for Cancer Research (AFCR17SC01) to MZ, and grants from National Institute of Health (CA196878, CA217642, GM51586, DEO15964) to KLG. MZ is a Kerry Holdings Professor in Science and a Senior Fellow of IAS at HKUST.

## Author contributions

ZL, ZY, and RX performed the experiments; ZL, ZY, RX, ZJ, KLG and MJZ analyzed data; ZL, ZY, and MZ wrote the paper and all authors approved the manuscript; MZ and KLG supervised the research; MZ coordinated the project.

## Supplementary Figures and Table

**Figure S1.**
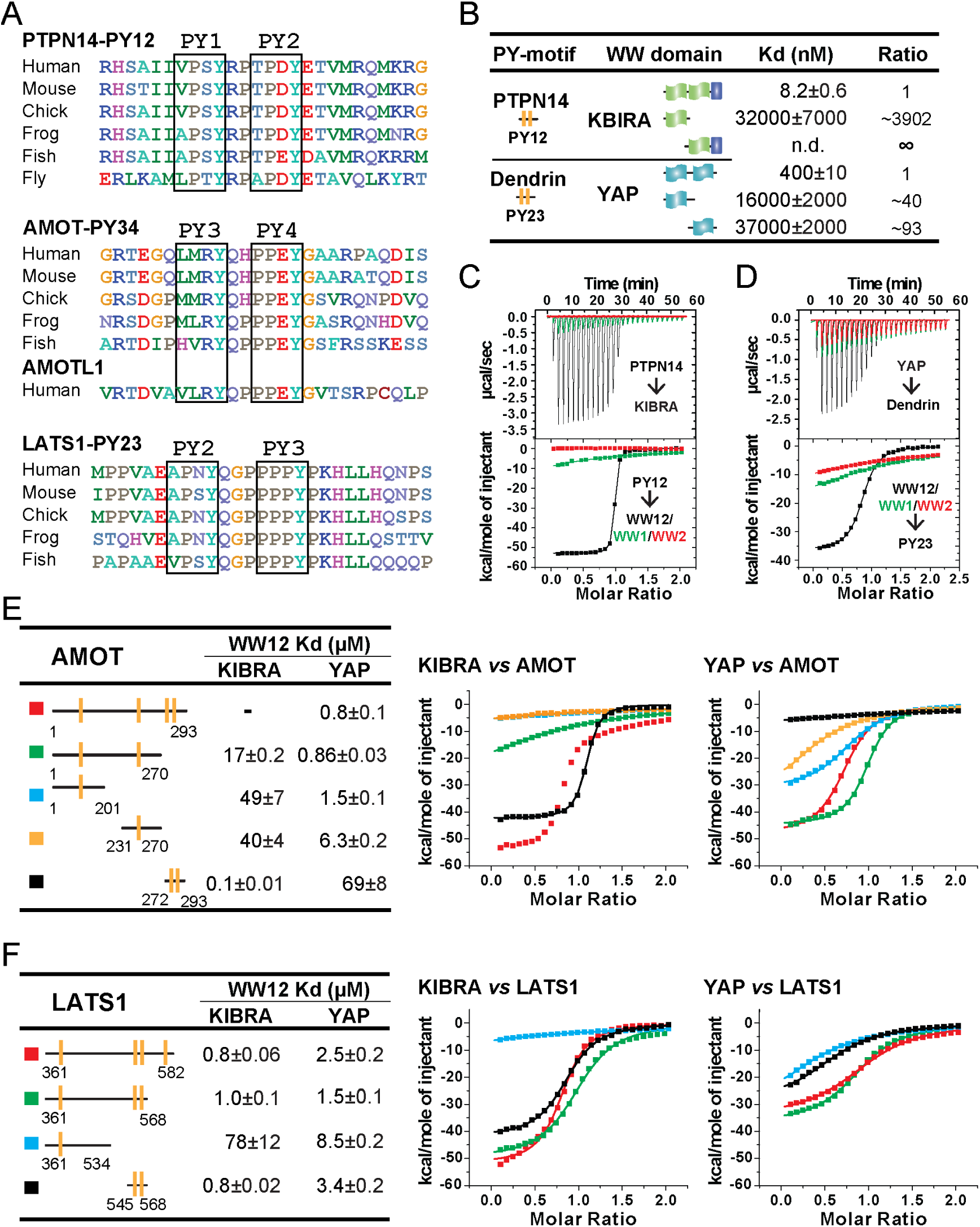
Super strong bindings of the KIBRA WW tandem to PY-motif containing proteins functioning in cell growth control. (A) Amino acid sequence alignment of the closely spaced PY motifs from PTPN14, AMOTs and LATS1. (B) ITC-derived binding affinities of isolated WWs and WW tandems from KIBRA and YAP in binding to closely connected PY motif. (C&D) Superimposition of ITC curves of isolated WWs and WW tandems from KIBRA (C) and YAP (D) in binding to closely spaced PY motif. (E&F) ITC-based measurements of the bindings between WW tandems from KIBRA and YAP, and various PY motifs of AMOT (E) and LATS1 (F).

**Figure S2.**
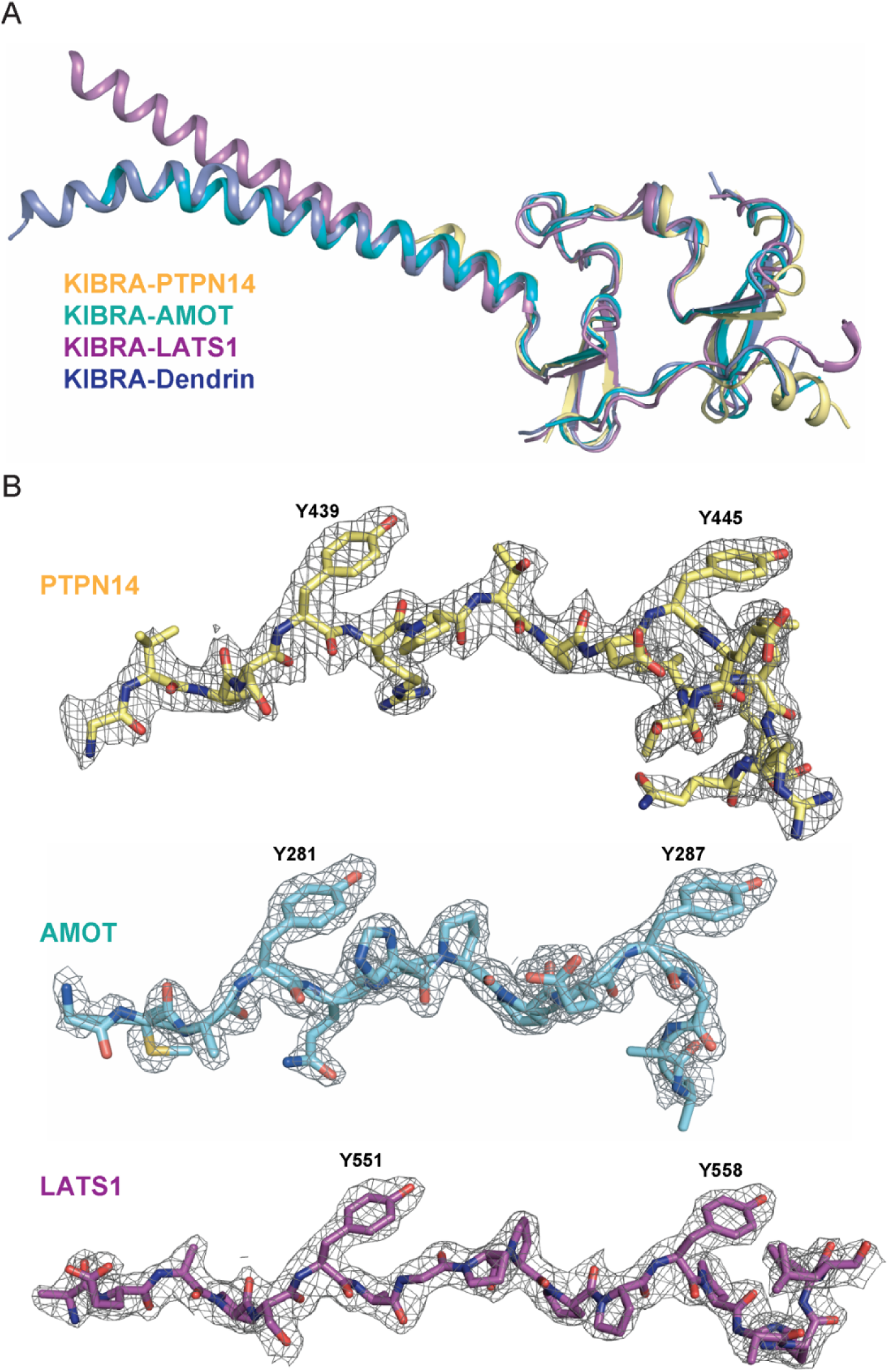
Structures of KIBRA WW12 in Complex with various closely connected PY motifs. (A) Superimposition of the overall structures of the KIBRA WW tandem in complex with PY motifs from PTPN14, AMOT, LATS1, and Dendrin. (B) The electron density maps of PY motifs from PTPN14, AMOT and LATS1 are shown and are colored grey and contoured at 1.5δ.

**Figure S3.**
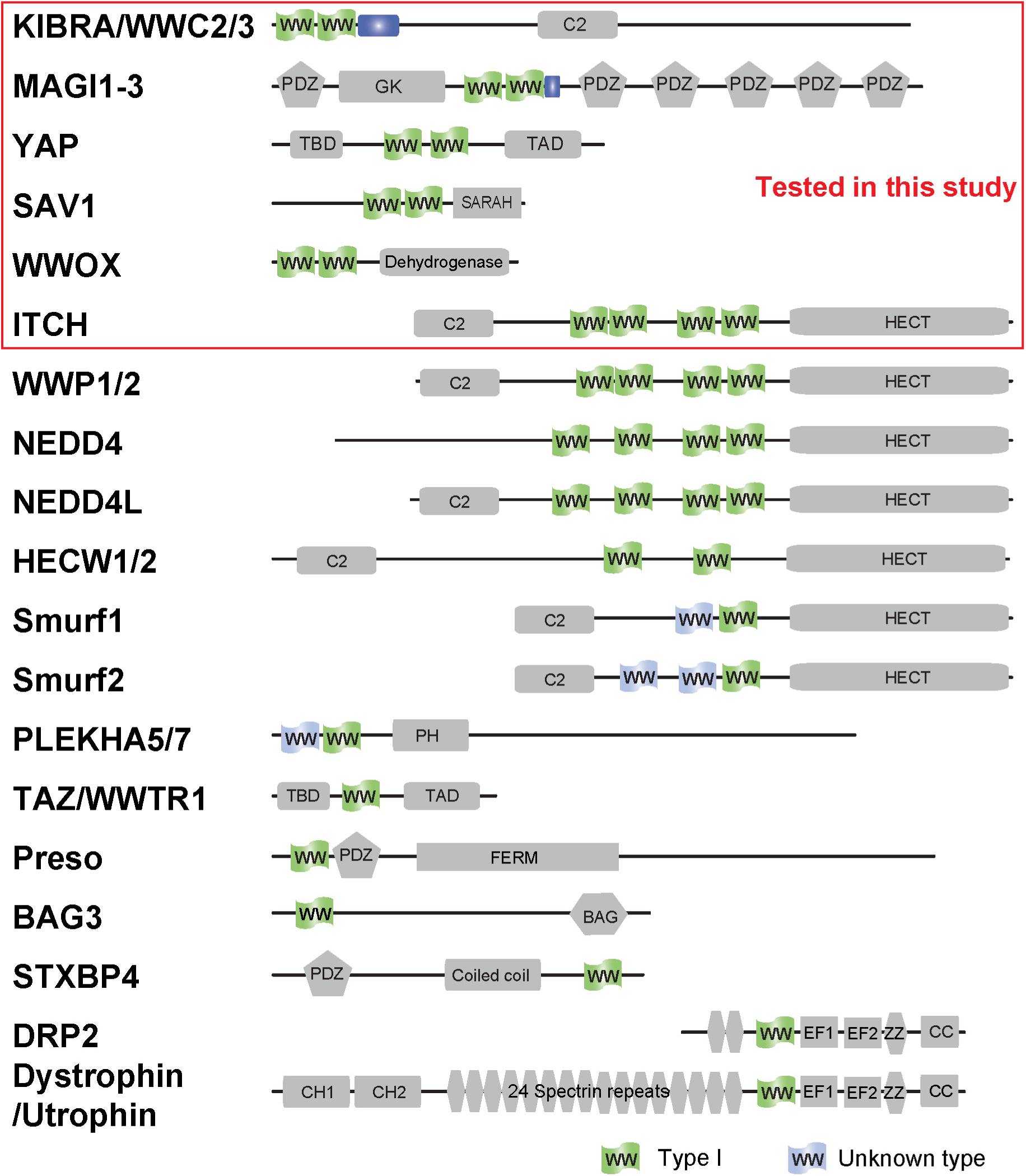
Domain organization of Type I WW containing human proteins.

**Figure S4.**
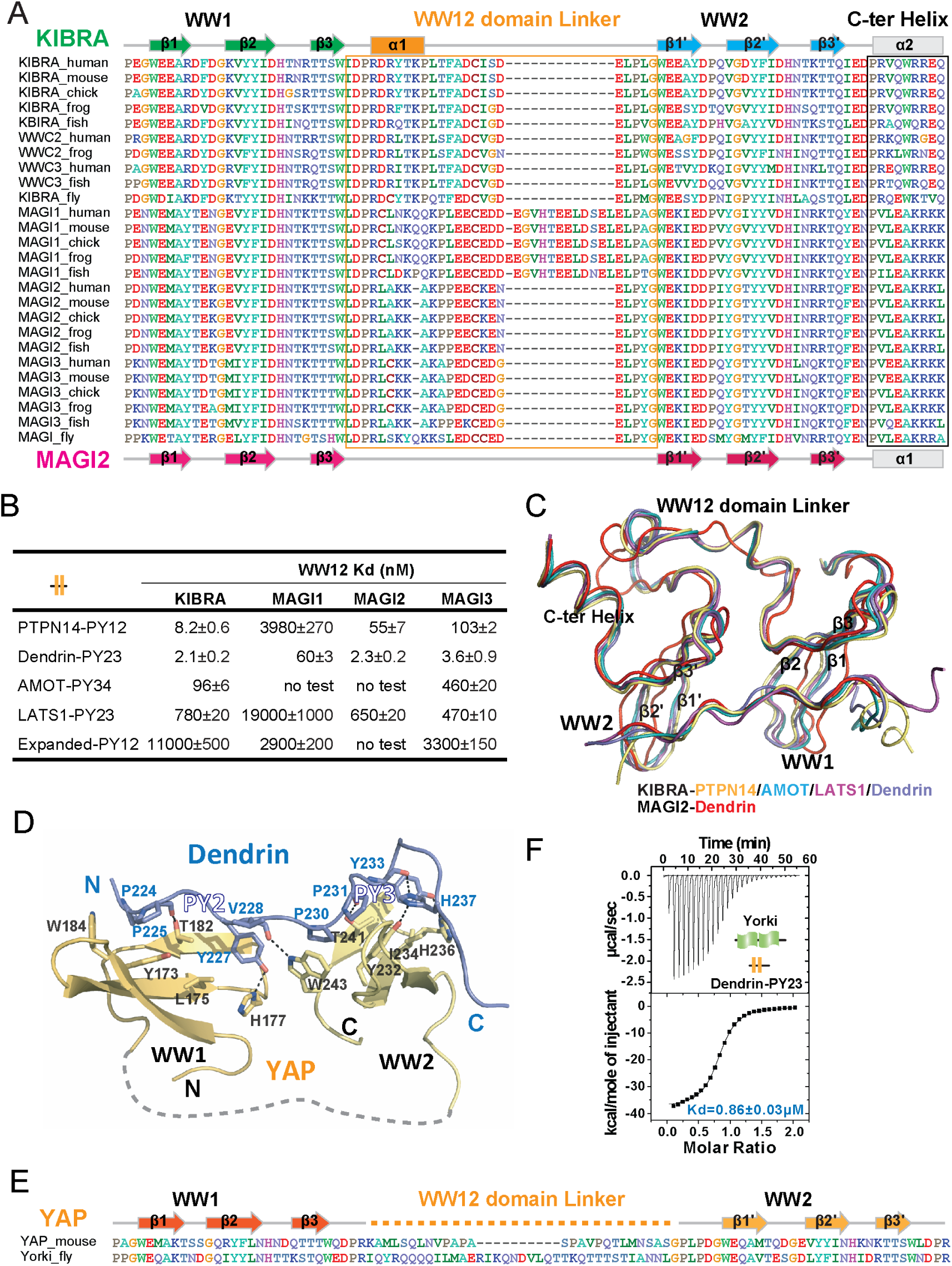
Structural basis governing the strong bindings of various WW tandems to their PY-motif containing targets. (A) Amino acid sequence alignment of the WW tandems of KIBRA/WWC and MAGI families from different species. (B) ITC-based measurements comparing the binding affinities between various closely spaced PY motifs and WW tandems from KIBRA and MAGI family. (C) Superimposition of the complex structures of KIBRA and MAGI2 WW tandems with their respective ligands. (D) Ribbon and stick model showing the detailed interactions of YAP WW tandem in complex with Dendrin-PY23. (E) Amino acid sequence alignment of the WW tandems of mouse YAP and *Drosophila* Yorkie. (F) ITC-based measurement of the binding between Yorkie WW tandem and Dendrin-PY23.

**Figure S5.**
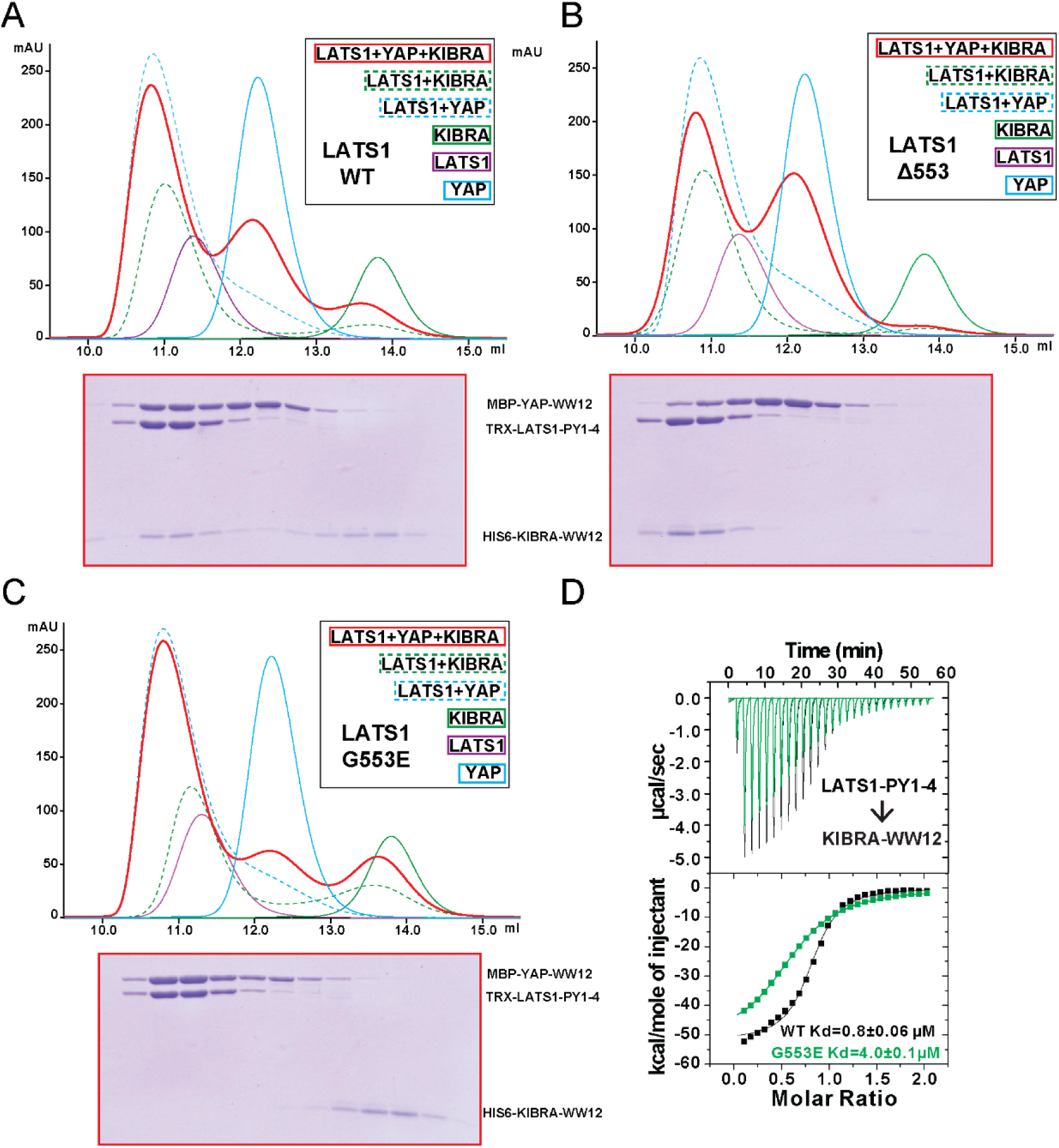
KIBRA and YAP compete for binding to LATS1. (A) Analytical gel filtration chromatography showing that KIBRA WW12 and YAP WW12 compete with each other for binding to WT LATS1 PY1-4. The elution fractions of the KIBRA WW12, YAP WW12 and LATS1 PY1-4 (or its mutants in Panels B&C below) mixture at a 1:1:1 ratio (red curve) were analyzed by SDS-PAGE. The proteins used were: His6-KIBRA WW12 (aa 5-97), MBP-tagged YAP WW12 (aa 151-251), and Trx-tagged LATS1 PY1-4 (aa 361-582) or the corresponding Gly553 mutants below. (B) LATS1 PY1-4(Δ553) displayed enhanced binding to KIBRA WW12, as all KIBRA WW12 was shift to the complex fractions and YAP WW12 shifted to the free form. (C) LATS1 PY1-4(G553E) cannot bind to KIBRA WW12 as all KIBRA WW12 shifted to the free form, and LATS1 PY1-4(G553E) became prefretially bound to YAP WW12. (D) ITC-based measurements of the bindings of KIBRA WW12 to WT LATS1 PY1-4 or LATS1 PY1-4(G553E).

**Figure S6.**
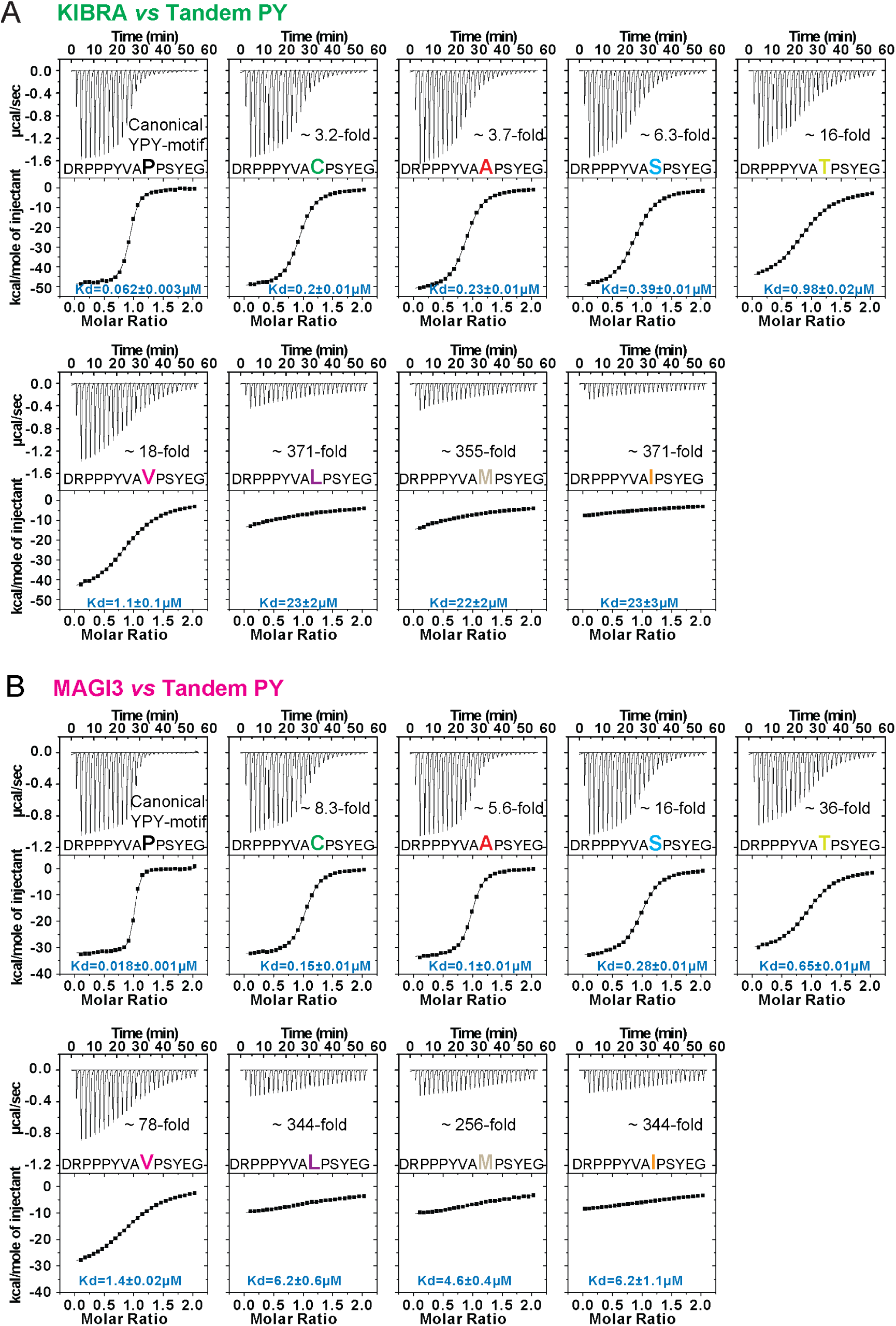
Bindings between WW tandems and closely spaced PY motifs with 2^nd^ PY variation. (A&B) ITC-based measurements comparing the binding of WW tandems from KIBRA (A) and MAGI3 (B) to Dendrin-PY23 with variations on 2^nd^ PY motif.

**Figure S7.**
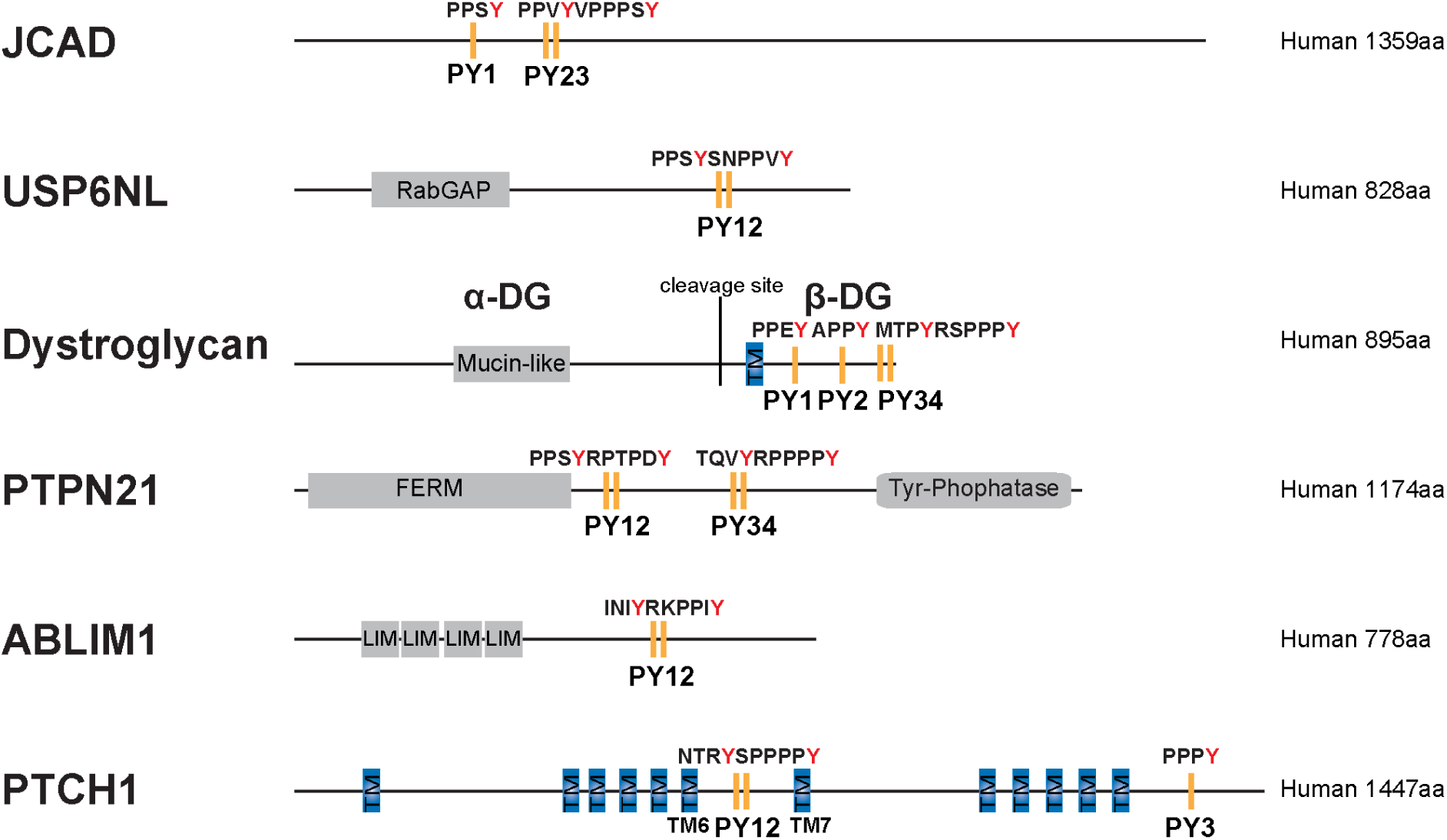
Domain organization of selected proteins with PY motifs fitting the consensus sequence indicated in panel Fig. 7A.

**Table S1:** Statistics of Data Collection and Model Refinement.

|  | **KIBRA/**<br>**Lats1-Se-Met** | **KIBRA/**<br>**LATS1** | **KIBRA/**<br>**PTPN14** | **KIBRA/**<br>**AMOT** | **KIBRA/**<br>**β-DG** | **MAGI2/**<br>**Dendrin** | **Dendrin-**<br>**“L”-YAP1** | **YAP1-“L”**<br>**-Dendrin** |
| --- | --- | --- | --- | --- | --- | --- | --- | --- |
| **Data collection** |  |  |  |  |  |  |  |  |
| Space group | P2<sub>1</sub>2<sub>1</sub>2<sub>1</sub> | P2<sub>1</sub>2<sub>1</sub>2<sub>1</sub> | R32 | P2<sub>1</sub>2<sub>1</sub>2 | P321 | P2<sub>1</sub>2<sub>1</sub>2<sub>1</sub> | P4<sub>1</sub>2<sub>1</sub>2 | P6<sub>3</sub>22 |
| Unit cell (Å) | 44.2,64.8,166.6 | 44.3,62.8,156.8 | 107.6,107.6,107.8 | 65.6,72.3,79.4 | 73.5,73.5,105.3 | 46.9,58.6,86.3 | 77.4,77.4,116.8 | 94.6,94.6,174.4 |
| Resolution (Å) | 50-3.10<br>(3.15-3.10) * | 50-2.50<br>(2.54-2.50) | 50-2.40<br>(2.44-2.40) | 50-2.0<br>(2.03-2.0) | 50-2.30<br>(2.34-2.30) | 50-1.65<br>(1.68-1.65) | 50-2.0<br>(2.03-2.0) | 30-3.10<br>(3.15-3.10) |
| Redundancy | 6.7(6.8) | 11.2(9.9) | 9.9(10.3) | 9.9(10.2) | 10.5(10.7) | 6.9(6.8) | 12.8(11.2) | 34.6(35.4) |
| I/σI | 30.0(2.2) | 23.7(2.1) | 18.2(2.7) | 22.4(2.2) | 40.0(2.7) | 44.0(2.5) | 32.8(3.7) | 15.3(2.0) |
| Completeness (%) | 97.5(98.9) | 99.5(91.1) | 100(100) | 99.8(99.9) | 99.7(100) | 99.8(99.9) | 100(100) | 99.9(100) |
| *R*<sub>sym</sub> or *R*<sub>merge</sub> | 0.055(0.802) | 0.102(0.982) | 0.124(0.957) | 0.103(0.752) | 0.056(0.886) | 0.047(0.856) | 0.079(0.698) | 0.225(>1) |
| No. reflections | 17,894 | 15,085 | 9,363 | 26,123 | 14,385 | 29,302 | 23,948 | 8,870 |
| **Refinement** |  |  |  |  |  |  |  |  |
| Resolution (Å) |  | 50-2.50 | 50-2.40 | 50-2.0 | 50-2.30 | 50-1.65 | 50-2.0 | 50-3.10 |
| *R*<sub>cryst</sub> / *R*<sub>free</sub> |  | 0.226/0.274 | 0.190/0.242 | 0.21/0.235 | 0.21/0.253 | 0.206/0.238 | 0.187/0.211 | 0.216/0.243 |
| No. of atoms |  |  |  |  |  |  |  |  |
| Protein/Water |  | 2,190/80 | 904/32 | 2,254/276 | 1,214/75 | 1,693/144 | 1,433/198 | 879/5 |
| B-factors |  |  |  |  |  |  |  |  |
| Protein/Water |  | 42.1/38.2 | 46.0/47.6 | 41.5/45.3 | 43.8/39.5 | 34.7/39.4 | 32.6/40.2 | 58.3/56.5 |
| R.m.s. deviations |  |  |  |  |  |  |  |  |
| Bond lengths (Å) |  | 0.011 | 0.008 | 0.008 | 0.009 | 0.006 | 0.007 | 0.018 |
| Bond angles (°) |  | 1.261 | 0.984 | 1.14 | 1.101 | 1.003 | 1.103 | 1.265 |
| Ramachandran plot |  |  |  |  |  |  |  |  |
| Favored/Allowed<br>/Outliers (%) |  | 95.5/4.5/0 | 100/0/0 | 99.3/0.7/0 | 97.9/2.1/0 | 99.4/0.6/0 | 99/1/0 | 94.4/5.6/0 |

\*Highest-resolution shell is shown in parentheses

